# An IL-1β driven neutrophil-stromal cell axis fosters a BAFF-rich microenvironment in multiple myeloma

**DOI:** 10.1101/2023.03.03.530773

**Authors:** Madelon M.E. de Jong, Cathelijne Fokkema, Natalie Papazian, Teddie van Heusden, Michael Vermeulen, Remco Hoogenboezem, Gregory van Beek, Sabrin Tahri, Mathijs A. Sanders, Pieter van de Woestijne, Francesca Gay, Philippe Moreau, Maike Büttner-Herold, Heiko Bruns, Mark van Duin, Annemiek Broijl, Pieter Sonneveld, Tom Cupedo

## Abstract

The bone marrow permanently harbors high numbers of neutrophils, and a tumor-supportive bias of these cells could significantly impact bone marrow-confined malignancies. In multiple myeloma, the bone marrow is characterized by inflammatory stromal cells with the potential to influence neutrophils. We investigated myeloma-associated alterations in marrow neutrophils and the impact of stromal inflammation on neutrophil function. Mature neutrophils in myeloma marrow are activated and tumor-supportive, transcribing increased levels of IL-1β, and myeloma cell survival factor BAFF. Interactions with inflammatory stromal cells can induce neutrophil activation, including BAFF secretion, in a STAT3-dependent manner and once activated, neutrophils gain the ability to reciprocally induce stromal activation. After first-line myeloid-depleting treatment, patient bone marrow retains residual stromal inflammation and newly-formed neutrophils are reactivated. Combined, we identify a neutrophil-stromal cell feed-forward loop driving tumor-supportive inflammation that persists after treatment and warrants novel strategies to target both stromal and immune microenvironments in multiple myeloma.

## Introduction

The tumor microenvironment (TME) plays a pivotal role in cancer pathobiology and is often characterized by local, subclinical inflammation that impacts tumor development, progression and therapy resistance^1^. Multiple myeloma (MM) is a hematological malignancy defined by the accumulation of malignant plasma cells in the bone marrow (BM)^2^, and interactions between myeloma cells and their BM environment are crucial for tumor survival^3^. Recently, we identified MM-specific inflammatory BM mesenchymal stromal cells (iMSC) that are induced by inflammatory cytokines such as interleukin (IL)-1β, and are a source of plasma cell survival factors^4^. Moreover, iMSC transcribe ligands for neutrophil-expressed C-X-C Motif Chemokine Receptors 1 (CXCR1) and CXCR2^5^, as well as other neutrophil modulation factors such as IL-6 and complement component 3 (C3)^4, 6, 7^.

The tumor microenvironment of solid tumors is often infiltrated by high numbers of mature neutrophils^8^, that play a crucial role in the establishment of cancer-associated inflammation^9^. Neutrophils can stimulate cancer cell proliferation, promote neoangiogenesis and suppress the cytotoxic anti-tumor immune response^10^. Consequently, high numbers of intra-tumoral neutrophils are associated with poor prognosis, increased metastasis, and advanced tumor stage^11, 12^. As the primary site of neutrophil development, the BM permanently harbors the highest number of neutrophils in the human body^13^. This raises the possibility that such preset abundance of neutrophils could impact tumor pathobiology in BM-confined malignancies such as MM.

Neutrophils from BM or blood of MM patients are immune suppressive^14,15,16^, limiting T-cell proliferation *in vitro*. Moreover, in newly-diagnosed patients, high BM mature neutrophil/T-cell ratios^15^ and blood neutrophil/lymphocyte ratios^17, 18^ are associated with inferior progression-free survival. Interestingly, MSC from patients with MM can induce the T-cell suppressive phenotype in neutrophils^19^. Tumor-supportive functions of neutrophils beyond immune suppression are less well defined in MM. Considering the presence of stromal inflammation in MM that may affect the BM neutrophil compartment, we hypothesized that stromal–neutrophil interactions might significantly impact the tumor-supportive TME in MM.

Here, we used single cell transcriptomics, flow cytometry and functional studies to identify inflammatory, tumor-supportive mature neutrophils in the BM of newly diagnosed MM patients. Activated neutrophils transcribed high levels of pro-inflammatory cytokines, including IL-1β, and myeloma cell survival factors, such as B cell maturation antigen (BCMA)-ligand B cell activating factor (BAFF). The tumor supportive phenotype of BM neutrophils, including BAFF secretion, was induced by inflammatory stroma in a STAT3-dependent manner. Conversely, activated neutrophils gained the ability to induce stromal inflammation via IL-1β, suggestive of pathogenic amplification of a tumor-supportive microenvironment. BAFF-transcribing neutrophils, elevated BAFF BM plasma protein levels and inflammatory stromal cells persisted after first-line treatment, even in the absence of detectable tumor burden. These findings reveal a stromal cell-neutrophil axis leading to a tumor-supportive TME with potential implications for MM cell survival and disease relapse.

## Results

### Single cell transcriptomic overview of the BM myeloid compartment

Mature neutrophils are the most abundant nucleated cell in BM aspirates of newly diagnosed MM (NDMM) patients, and 12-35x more prevalent than any other myeloid subset (37.7 ± 9.0% mature neutrophils vs. 1.4 ± 0.4% macrophages and 0.8 ± 0.6% monocytes, Figure 1A, Supplemental Figures 1A-B). To identify neutrophil alterations in MM, we generated a single cell transcriptomic overview of the entire myeloid lineage from fresh BM aspirates of 6 NDMM patients and 4 age-matched non-cancer controls (Supplemental Tables 1 and 2). Briefly, all CD45-positive cells contained within granulocytic and monocytic scatter populations were sort-purified, and presence of neutrophils and monocytes/macrophages was verified by expression of CD11b/CD16 and CD14/CD163, respectively (Supplemental Figures 1A-B). Single cell sequencing led to the generation of a dataset of 90,168 myeloid cells (52,542 of NDMM BM and 37,626 of control BM, Supplemental Figure 1C). This dataset could broadly be divided in three major populations: myeloid progenitors (*MPO*^+^), mononuclear phagocytes (*CD14*^+^), and neutrophils (*ELANE*^+^ or *MME*^+^, Supplemental Figures 1C-D). Myeloid progenitors consisted of *MPO*^+^*ELANE*^+^ myeloblasts, *CD14*^+^*MPO*^+^ monoblasts, and a cluster of various cycling progenitors (*MKI67*^+^, Supplemental Figure 1E). The mononuclear phagocyte population contained *CD14*^+^*FCGR3A*^—^ classical and *CD14*^—^*FCGR3A*^+^ non-classical monocytes, *CD163*^+^*IL1B*^+^ pro-and *CD163*^+^*LTF*^+^ anti-inflammatory macrophages and *CD1C*^+^ dendritic cells (Supplemental Figure 1E). Importantly, the entire neutrophilic lineage was captured, starting from *ELANE*^+^*AZU1*^+^ myelocytes, via *LTF*^+^ pre-neutrophils (PreNeu), *ARG1*^+^*MMP9*^+^ immature neutrophils (ImmNeu) into *MME*^+^ mature neutrophils (MatNeu, Supplemental Figure 1E). Developmental trajectories and the neutrophil-monocyte lineage branching point were confirmed by pseudotime analysis (Supplemental Figure 1F). Quantification of cluster abundance did not show significant differences between NDMM BM and controls (Supplemental Figures 1G-H). The entire dataset is available for interactive browsing at www.bmbrowser.org.

**Figure 1.**
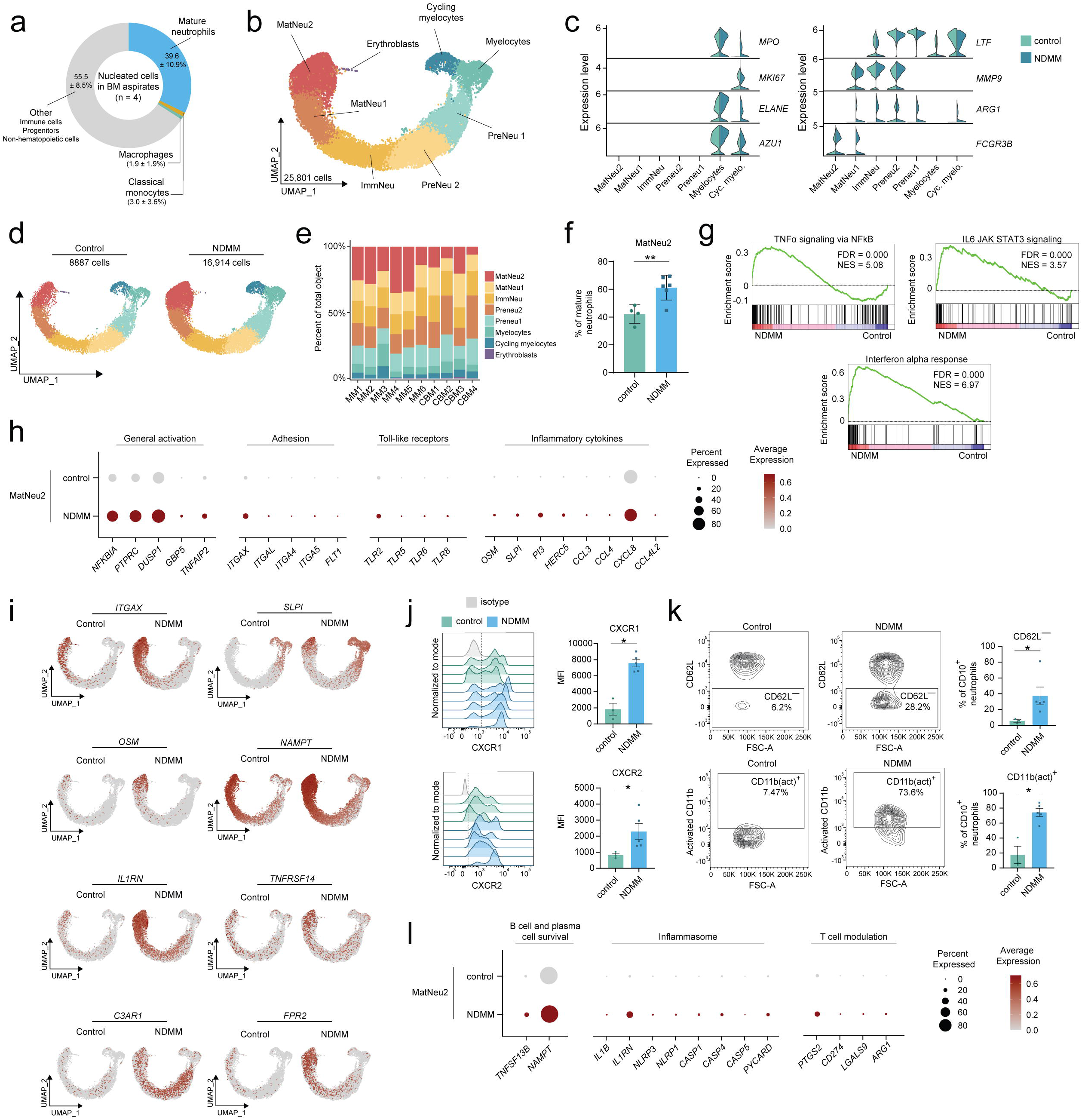
NDMM BM is enriched for inflammatory mature neutrophils. **A)** Distribution of nucleated cells in BM aspirates of n = 4 NDMM patients, values are mean ± SD **B)** UMAP of the combined dataset of 16,914 cells from 6 NDMM patients and 8,887 cells from 4 non-cancer controls showing 7 clusters identified by marker genes in *(C)*. Colors represent clusters; UMAP, uniform manifold approximation and projection **C)** Marker gene transcription in the 7 clusters, split by condition (light green, controls; dark green, NDMM patients) **D)** Original UMAP split into non-cancer control and NDMM datasets **E)** Cluster distribution per patient. Colors represent clusters. ‘MM’, MM patient. ‘CBM’, control **F)** Percentages of mature neutrophils in cluster MatNeu2 per individual in each condition **G)** Enrichment plots for selected Hallmark gene sets in cluster MatNeu2 comparing NDMM patients to controls; FDR, false discovery rate; NES, normalized enrichment score **H)** Transcription of differentially expressed genes in cluster MatNeu2 comparing controls to NDMM patients. Size of dot represents fraction of cells transcribing the gene, color represents level of transcription **I)** Transcription of selected differentially expressed genes in control and NDMM datasets **J)** Representative histogram of expression of CXCR1 and CXCR2 on CD15^+^CD16^+^ mature neutrophils of non-cancer controls (green) and NDMM patients (blue) measured by flow cytometry. Isotype controls are depicted in grey. Mean fluorescent intensity (MFI) of CXCR1/2 was quantified for each individual per condition **K)** Representative flow plots of expression of CD62L and the activated conformation of CD11b (‘CD11b(act)’) on CD10^+^ mature neutrophils of non-cancer controls and NDMM patients. Frequencies of CD62L^—^ and CD11b(act)^+^ cells were quantified for each individual per condition. **L)** Transcription of differentially expressed genes in cluster MatNeu2 comparing controls to NDMM patients. Size of dot represents fraction of cells transcribing the gene, color represents level of transcription Data are presented as mean ± SEM. Significance was calculated in **F**, **J** and **K** using the Mann– Whitney U test (two-tailed), *P ≤ 0.05, **P ≤ 0.01.

### NDMM BM is enriched for inflammatory mature neutrophils

To allow detailed investigation of MM-associated alterations in neutrophils, the neutrophilic lineage was reintegrated to form a new dataset in which clustering was only driven by neutrophil-derived transcripts (see methods). The major subpopulations in the neutrophil dataset largely mirrored those in the myeloid lineage dataset, with the exception of myelocytes, which now segregated in a cycling (*MKI67*^+^) and non-cycling cluster (Figures 1B-C) and mature neutrophils that now segregated in two separate clusters (MatNeu1 and MatNeu2, Figure 1B-C). Interestingly, cluster MatNeu2 was significantly enriched in NDMM BM compared to controls (Figures 1D-F, Supplemental Figure 1I).

In MM patients, cluster MatNeu2 was defined by inflammatory gene programs, including enrichment of the NFκB-, interferon- and STAT3 pathway gene sets (Figure 1G, Supplemental Table 3). Fittingly, differentially expressed genes (DEGs) of the MatNeu2 cluster in NDMM compared to non-cancer controls were mostly related to inflammatory activation of neutrophils (*PTPRC*, *DUSP1*, *GBP5*), adhesion and complement sensing (*ITGAX*, *ITGAL, ITGA4*), innate sensing (various TLRs), and inflammatory mediators produced by activated neutrophils (*OSM*, *CXCL8*, *CXCL16*, Figures 1H-I, Supplemental Table 4). A subset of neutrophils in MatNeu2 was defined by interferon (IFN) response genes (Supplemental Figure 1J). Cellular activation and inflammation were also the factors distinguishing neutrophils in MatNeu2 from those in cluster MatNeu1 (Supplemental Figures 1K and 2A, Supplemental Tables 5 and 6). To validate mature neutrophil activation in NDMM BM, we analyzed BM cells from 5 NDMM patients and 3 non-cancer controls by flow cytometry. Mature neutrophils of patients with MM showed increased expression of CXCR1 and CXCR2 (Figure 1J, Supplemental Figure 2B-D), loss of CD62L (Figure 1K, Supplemental Figure 2B-C) and a switch to the activated configuration of integrin CD11b compared to controls (Figure 1K, Supplemental Figure 2B-C). Combined, these findings identify mature BM neutrophils in NDMM as activated and pro-inflammatory.

**Figure 2.**
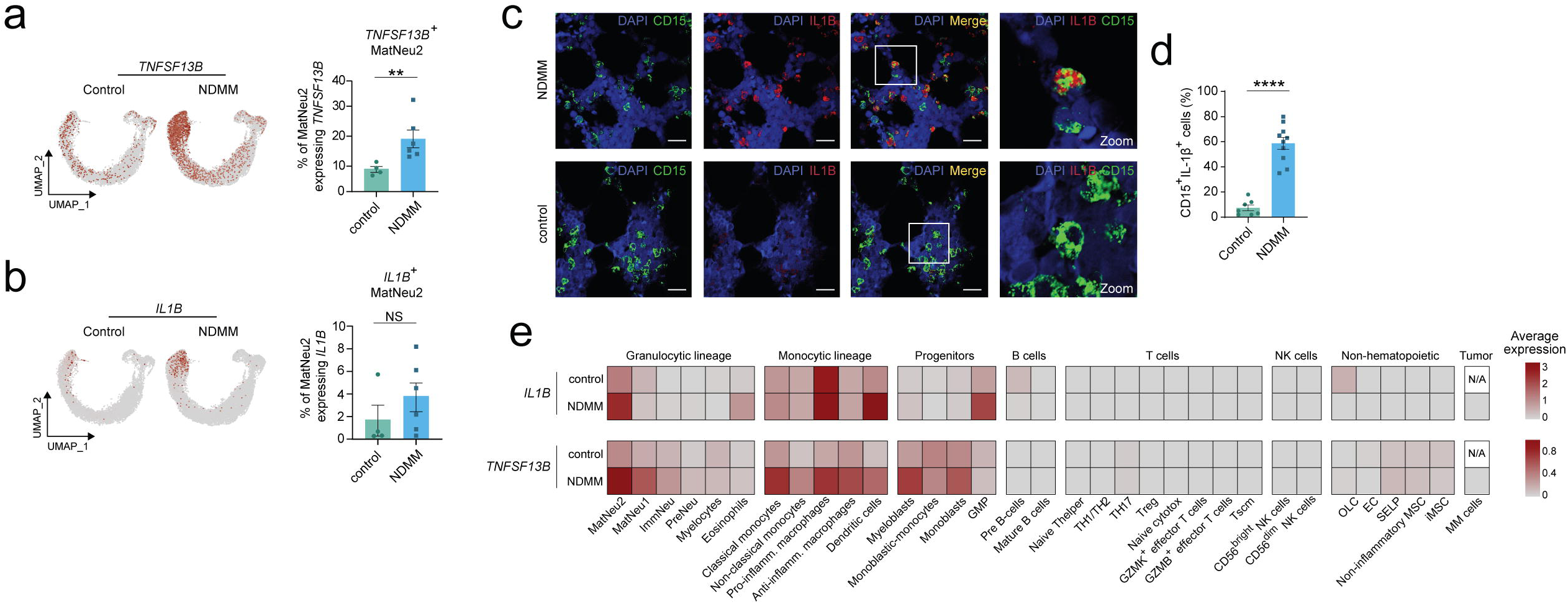
Mature neutrophils are a considerable source of IL1B and TNFSF13B. **A)** Transcription of *TNFSF13B* in non-cancer controls and NDMM patients. Frequencies of *TNFSF13B*-transcribing cells in the MatNeu2 cluster were quantified for each individual per condition **B)** Transcription of *IL1B* in non-cancer controls and NDMM patients. Frequencies of *IL1B*-transcribing cells in the MatNeu2 cluster were quantified for each individual per condition **C)** Representative images of BM biopsies of N = 10 NDMM patients (upper panel) and N = 7 non-cancer controls (lower panel) stained for CD15 (green), IL-1β (red), and DNA (DAPI, blue). Pictures were taken at a magnification of 630x. Scale bar: 20 μm **D)** Quantification of CD15^+^IL-1β^+^ cells in 25 regions of interest (ROIs) per individual in each condition **E)** Average transcription of *IL1B* and *TNFSF13B* per annotated cluster from single cell RNA sequencing datasets. Data is split in average expression in non-cancer controls and NDMM patients. Data are presented as mean ± SEM. Significance in **A**, **B** and **D** was calculated using the Mann–Whitney U test (two-tailed); **P ≤ 0.01, ****P ≤ 0.0001, NS P > 0.05.

Neutrophils within cluster MatNeu2 of NDMM patients also displayed significantly increased transcription of genes associated with MM or plasma cell biology. Such genes included nicotinamide phosphoribosyltransferase (*NAMPT*) (Figures 1I, L), a neutrophilic survival factor^20^ that also functions as a cytokine promoting B cell maturation^21^ and myeloma cell proliferation^22^, and that is elevated in blood plasma of MM patients^23^. Although generally recognized as an adipokine, our data suggest that mature neutrophils may also be a source of *NAMPT* in MM. Moreover, cells in MatNeu2 transcribed high levels of *TNFSF13B*, which encodes the BCMA-ligand BAFF^24^ (Figure 1L). BCMA is ubiquitously expressed by myeloma cells^25^ and ligation with BAFF increases growth of myeloma cells in culture^26^ (Supplemental Figure 2E). Increased transcription of *TNFSF13B* in MM bone marrow is also associated with rapid disease progression^27^.

Mature neutrophils also showed signs of inflammasome priming, a prerequisite for secretion of IL-1β^28, 29^, a pro-inflammatory cytokine associated with MM disease progression^30, 31^ and elevated in patient BM plasma^4^. Transcripts of canonical (*NLRP3*, *CASP1*) and non-canonical inflammasome components (*NLRP1*, *CASP4*, *CASP5*) were elevated, as was transcription of *IL1B* and the co-transcribed antagonist *IL1RN* (Figures 1I, L). Finally, in line with the reported immune-modulatory function of MM-associated neutrophils^15^, a small percentage of cells in the MatNeu2 cluster transcribed genes associated with T cell inhibition, such as *PTGS2* (encoding COX2) ^32^ and ligands for inhibitory checkpoint molecules, including *CD274* (PD-L1), *LGALS9* (galectin-9) and *TNFRSF14* (HVEM, Figures 1I, L). Together, these data indicate that mature neutrophils in MM BM are characterized by tumor-supportive and immune-modulatory transcripts.

### Mature neutrophils in NDMM are a considerable source of IL1B and TNFSF13B

*IL1B* and *TNFSF13B* are MM pathobiology-related genes^30, 31, 33, 34^ with increased transcription in NDMM neutrophils compared to controls (Figure 1H, Figures 2A, B). The percentage of mature neutrophils transcribing *TNFSF13B* was also significantly increased in NDMM BM (Figure 2A). The percentages of *IL1B*^+^ mature neutrophils displayed large interpatient variation at the transcript level (Figure 2B). Nevertheless, enumeration of IL-1β-expressing CD15^+^ neutrophils in BM biopsies of NDMM patients revealed a significant increase compared to controls (Figures 2C-D).

We sought to investigate the extent of neutrophilic contribution to the overall transcription of *IL1B* and *TNFSF13B* and used our previously generated single cell sequencing datasets of the immune and non-hematopoietic TME^4^, together with the myeloid dataset described in the current study to analyze transcription of *TNFSF13B* and *IL1B* across the MM landscape. As expected, both genes were almost exclusively transcribed by myeloid populations (Figure 2E). MatNeu2 demonstrated one of the highest levels of transcription of *IL1B* in NDMM (Figure 2E). Transcription of *IL1B* was also high in pro-inflammatory macrophages, yet levels and percentages did not differ between NDMM patients and controls (Figure 2E, Supplemental Figures 2F-I). This notwithstanding, pro-inflammatory macrophages were enriched in the MM BM (Supplemental Figure 2J), and these cells may contribute to the increased IL-1β levels in MM BM plasma^4^. Increased transcription of *IL1B* was also observed in dendritic cells and granulocyte-monocyte progenitors (GMP), but both cell types are present in low frequencies in BM^35^, and percentages of positive cells did not differ between patients and controls (Supplemental Figures 2K-N).

*TNFSF13B* was increased in transcription across most myeloid populations of NDMM patients (Figure 2E). Inflammatory mature neutrophils (MatNeu2) displayed highest transcription of *TNFSF13B*, but non-inflammatory mature neutrophils (MatNeu1), classical monocytes, and macrophages all transcribed elevated levels of *TNFSF13B* (Figure 2E, Supplemental Figure 2O). However, only in MatNeu2 this increase in transcription was accompanied by an increase in percentage of *TNFSF13B*-transcribing cells (Figure 2A, Supplemental Figure 2P). Combined with the abundance of mature neutrophils in BM (Figure 1A), this suggests that activated mature neutrophils are a significant source of IL-1β and BAFF in NDMM.

### Inflammatory stromal cells induce the MM neutrophil phenotype

The MM BM is characterized by the presence of iMSC, which transcribe genes associated with neutrophil modulation. Therefore, we hypothesized that iMSC are involved in the induction of pro-inflammatory neutrophils in the myeloma BM. To this end, we developed an *in vitro* co-culture model to mimic iMSC – neutrophil interactions. As a stromal cell source, we used commercially-available adipose-derived stromal cells (ADSCs), which are often used to model MSC due to their tri-lineage differentiation potential and high degree of reproducibility^36^. Overnight stimulation of ADSC with recombinant IL-1β induced an iMSC-like transcriptome (Figure 3A-B, Supplemental Table 7), similar to stimulation of primary BM MSC^4^. Naïve peripheral blood neutrophils from healthy donors were subsequently co-cultured overnight with iMSC-like cells or unstimulated ADSC (referred to as MSC). Co-cultures with iMSC-like cells, but not non-inflammatory MSCs, induced the activated neutrophil expression profile, including upregulation of C3AR, CD11c, CD66b, CXCR1, CD45, and increased conformation of CD11b to its activated form (Figure 3C, Supplemental Figure 3A). Activation was not due to residual IL-1β from iMSC induction, as neutrophil cultures with recombinant IL-1β did not adopt this activated surface expression profile (Supplemental Figure 3B). Importantly, neutrophil activation was stromal cell-specific, as co-culture of neutrophils with MM cells failed to increase neutrophil activation markers, with the exception of CXCR1 (Supplemental Figure 3C). To analyze the extent of neutrophilic activation induced by iMSC-like cells, we analyzed iMSC-exposed neutrophils by RNA sequencing. Culture on MSC had a modest effect on neutrophils, leading to 89 differentially expressed genes (Supplemental Table 8). However, iMSC-exposed neutrophils increased transcription of 742 genes compared to neutrophils cultured alone and 481 genes compared to MSC-exposed neutrophils (Figure 3D, Supplemental Figure 3D, Supplemental Table 9). In line with inflammatory signaling in neutrophils from NDMM patients, iMSC-like cells increased NFκB-, and STAT3 signaling in neutrophils (Figure 3E), while the interferon signaling pathway that was present in NDMM patients was not induced by iMSC (Figure 3F). Many of the genes upregulated in iMSC-exposed neutrophils fitted into gene modules associated with inflammatory mature neutrophils from MM patients, including general activation genes, adhesion molecules, innate sensors, inflammatory cytokines, and components of the inflammasome including *IL1B* (Figures 3D, G). Co-culture with iMSC-like cells also induced neutrophilic transcription of the plasma cell survival gene *NAMPT*, while transcription of *TNFSF13B* showed considerable inter-donor variation (Figure 3G). Finally, neutrophil transcription of genes associated with modulation of the T cell response were unaffected by iMSC-like co-culture (Figure 3G), suggesting that induction of a T cell inhibitory phenotype in mature BM neutrophils is not the result of iMSC-signaling. Overall, the exposure of naïve neutrophils to activated stromal cells was sufficient to induce an inflammatory transcriptome highly similar to that of activated mature neutrophils in MM BM (Figure 3H, Supplemental Figure 3E).

**Figure 3.**
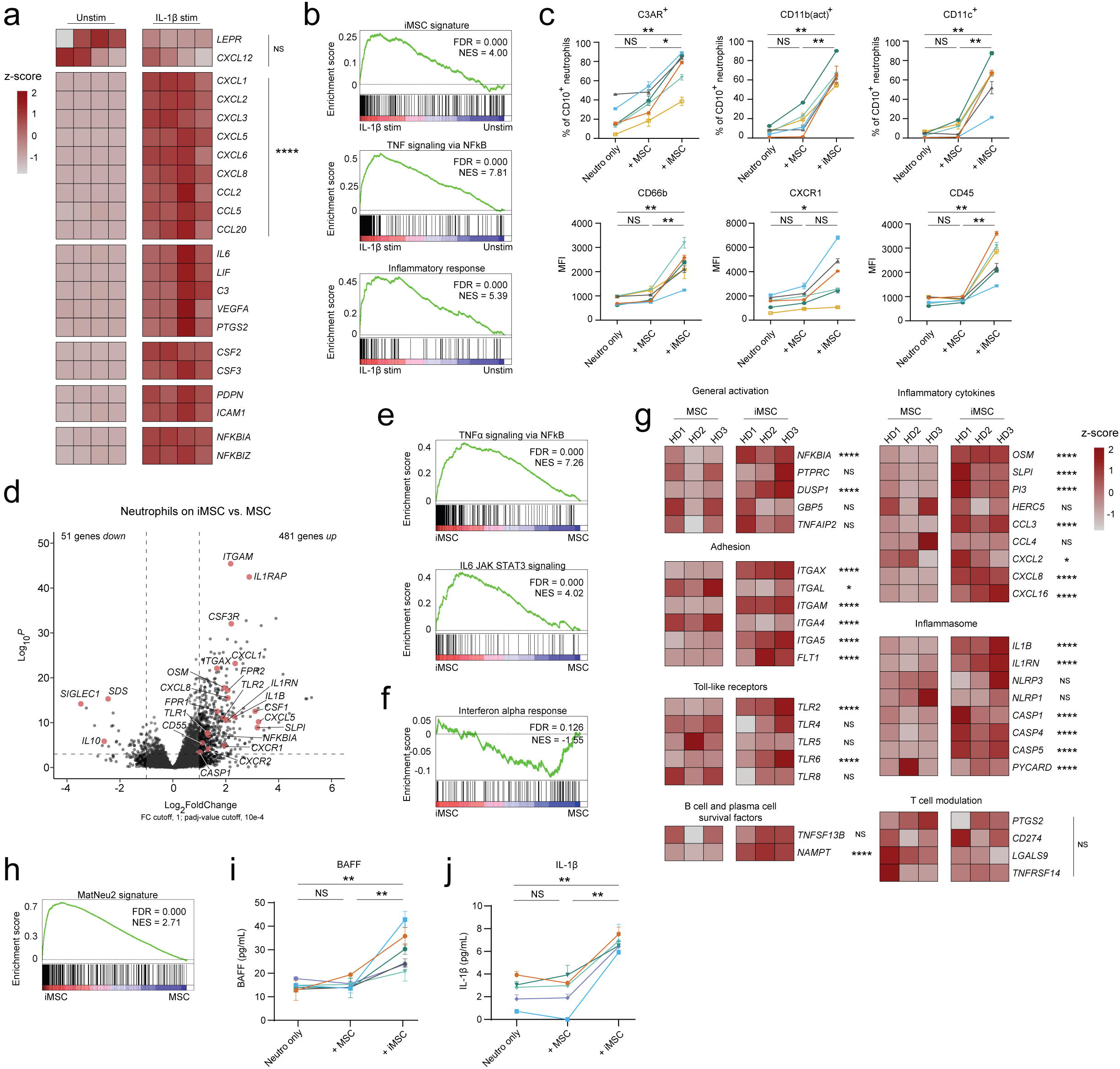
Inflammatory stromal cells induce the MM neutrophil phenotype. **A)** Transcription of inflammatory genes in IL-1β-stimulated and unstimulated ADSC (n = 4, collected over 4 separate experiments) **B)** Enrichment plots for the iMSC-signature (generated from DEGs of single cell RNA sequencing results in ^4^), and selected Hallmark genesets comparing stimulated and unstimulated ADSC **C)** Frequencies of CD10^+^ neutrophils expressing C3AR, the active conformation of CD11b (‘CD11b(act)’), and CD11c, or mean fluorescent intensity (MFI) of CD66b, CXCR1 and CD45 on CD10^+^ neutrophils cultured alone (‘neutro only’), on non-inflammatory stroma (‘+ MSC’), or on iMSC. Lines depict paired samples (n = 6, collected over 3 experiments) **D)** Volcano plot depicting differentially expressed genes of neutrophils cultured on iMSC versus neutrophils cultured on MSC. Log_2_FoldChange cutoff, 1; adjusted p-value (padj) cutoff 10^-4^. Genes in red are related to inflammation **E, F)** Enrichment plots for selected Hallmark gene sets comparing neutrophils cultured on iMSC to those cultured on MSC **G)** Transcription of MM-associated pro-inflammatory neutrophil gene modules in neutrophils cultured on MSC or on iMSC (n = 3 donors, collected in single experiment). HD, healthy donor **H)** Enrichment plot for the MatNeu2-signature (generated from DEGs of cluster MatNeu2 in NDMM patients) comparing neutrophils cultured on iMSC to those cultured on MSC **I)** BAFF or **J)** IL-1β protein in supernatant of neutrophils cultured alone (‘neutro only’), on non-inflammatory stroma (‘+ MSC’), or on iMSC. Lines depict paired samples (n = 6, collected over 5 experiments). Data are presented as mean ± SEM. Significance was calculated in **A, D** and **G** using the Wald test (two-tailed) followed by a Benjamini–Hochberg correction, and in **C, I** and **J** using the Wilcoxon Rank Sum test (two-tailed); *P ≤ 0.05, **P ≤ 0.01, ****P ≤ 0.0001, NS P > 0.05.

Analyses of protein secretion by neutrophils upon iMSC exposure revealed that, in contrast to transcription, protein secretion of BAFF was significantly increased (Figure 3I). Importantly, BAFF was not produced by iMSC-like cells (Supplemental Figure 3F) and was not induced by co-culture of neutrophils with myeloma cells (Supplemental Figure 3G). Protein levels of IL-1β were also increased after co-culture with iMSC-like cells compared to those cultured with non-inflammatory stroma or neutrophils alone (Figure 3J). This was not residual IL-1β from MSC activation, as iMSC conditioned medium did not contain increased levels of IL-1β (Supplemental Figure 3H). Increase of IL-1β secretion was stromal cell specific, as co-culture of neutrophils with myeloma cells did not induce IL-1β production (Supplemental Figure 3I). Together, these data indicate that inflammatory stromal cells can activate mature neutrophils and induce a pro-tumorphenotype, including production of BAFF and IL-1β.

### Neutrophil activation is STAT3-dependent

The inflammatory neutrophil transcriptome in MM patients was enriched for the STAT3-signaling gene set (Figure 1I), and patients displayed increased transcription of *STAT3* and STAT3 target gene *BCL3* (but not *SOCS3*) compared to controls (Figure 4A). *In vitro*, neutrophils co-cultured with iMSC-like cells increased transcription of *SOCS3*, *STAT3* and *BCL3* (Figure 4B), and increased intracellular levels of phosphorylated STAT3 protein (Figure 4C). To test the role of STAT3 signaling in the activation of neutrophils by iMSCs, we co-cultured naïve neutrophils with MSC or iMSC-like cells in the presence of the STAT3 inhibitor Stattic^37^, which inhibited *SOCS3* transcription (Supplemental Figure 3J), but did not affect neutrophil viability (Supplemental Figure 3K) or basic function (defined by expression of membrane metallo-proteinase CD10, Supplemental Figure 3L). However, STAT3 inhibition during co-culture with iMSC-like cells prevented neutrophil activation (Figure 4D) and BAFF secretion (Figure 4E), signifying the importance of this pathway. IL-1β release is generally regulated outside of STAT3, and consequently STAT3 inhibition did not affect neutrophil IL-1β secretion, indicating that neutrophil activation in MM is achieved by integration of multiple signals (Figure 4F).

**Figure 4.**
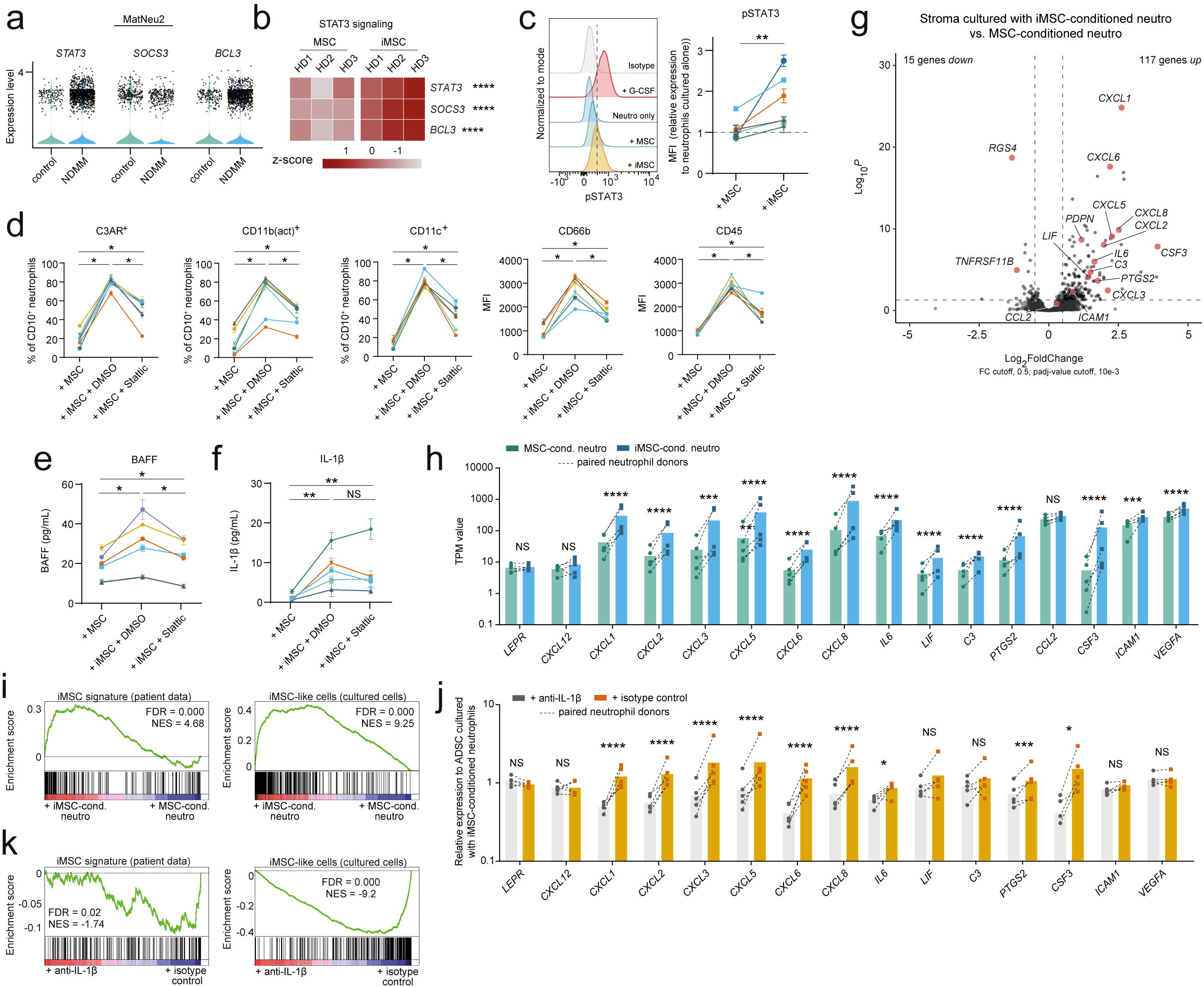
Neutrophil inflammation is induced by STAT3 signaling and inflammatory neutrophils can reciprocally induce the iMSC transcriptome. **A)** Transcription of STAT3, SOCS3 and BCL3 in cells in cluster MatNeu2 comparing control individuals and NDMM patients **B)** Transcription of genes associated with STAT3-signaling in neutrophils cultured on MSC or on iMSC (n = 3 donors, collected in single experiment). HD, healthy donor **C)** Representative histogram of expression of phosphorylated STAT3 (pSTAT3) from CD10^+^ neutrophils cultured in the presence of G-CSF (positive control), cultured alone, cultured on non-inflammatory stroma (‘MSC’), or cultured on iMSC, as determined by flow cytometry. Fold change in mean fluorescent intensity (MFI) compared to neutrophils cultured alone (dotted line) was quantified per individual of each condition. Lines depict paired samples (n = 6, collected over 6 experiments) **D)** Frequencies of CD10^+^ neutrophils expressing C3AR, CD11b(act), and CD11c, or mean fluorescent intensity (MFI) of CD66b and CD45 on CD10^+^ neutrophils cultured on non-inflammatory stroma (‘+ MSC’) or on iMSC in the presence of DMSO or Stattic. Lines depict paired samples (n = 6, collected over 3 experiments) **E)** BAFF protein in supernatant of neutrophils cultured on non-inflammatory stroma (‘+ MSC’) or on iMSC in the presence of DMSO or Stattic. Lines depict paired samples (n = 6, collected over 3 experiments). **F)** IL-1β protein in supernatant of neutrophils cultured on non-inflammatory stroma (‘+ MSC’) or on iMSC in the presence of DMSO or Stattic. Lines depict paired samples (n = 5, collected over 3 experiments) **G)** Volcano plot depicting differentially expressed genes of MSC cultured with iMSC-conditioned neutrophils versus MSC cultured with MSC-conditioned neutrophils. Log_2_FoldChange cutoff, 1; adjusted p-value (padj) cutoff 10^-3^. Genes in red are related to iMSCs. (n = 5, collected over 3 experiments) **H)** Transcription (transcripts per million, TPM) of iMSC-related genes in MSC cultured with MSC-conditioned neutrophils (green) and MSC cultured with iMSC-conditioned neutrophils (blue). Dotted lined depict paired samples. (n = 5, collected over 3 experiments) **I)** Enrichment plots for the iMSC-signature (generated from DEGs of single cell RNA sequencing results in ^4^), and iMSC-like cells signature (generated from data in Figure 5A) comparing MSC cultured with iMSC-conditioned neutrophils and MSC-conditioned neutrophils **J)** Transcription of iMSC-related genes in MSC cultured with iMSC-conditioned neutrophils in the presence of anti-IL-1β (grey) and in the presence of an isotype control (orange). Dotted lined depict paired samples (n = 5, collected over 3 experiments) **K)** Enrichment plots for the iMSC-signature (generated from DEGs of single cell RNA sequencing results in ^4^), and iMSC-like cells signature (generated from data in Figure 5A) comparing MSC cultured with iMSC-conditioned neutrophils in the presence of anti-IL-1β or in the presence of an isotype control. Data are presented as mean ± SEM. Significance was calculated in **B**, **G**, **H** and **J** using the Wald test (two-tailed) followed by a Benjamini– Hochberg correction, and in **C**, **D**, **E** and **F** using the Wilcoxon Rank Sum test (two-tailed); *P ≤ 0.05, **P ≤ 0.01, ***P ≤ 0.001, ****P ≤ 0.0001, NS P > 0.05.

### Inflammatory neutrophils are sufficient to induce the iMSC transcriptome

In contrast to MSC, neutrophils are motile cells that, in theory, could spread stromal inflammation within and between bones. We therefore tested whether iMSC-activated neutrophils can gain the ability to reciprocally activate non-inflammatory stromal cells. Neutrophils were co-cultured with iMSC-like cells to induce neutrophil activation and cells and supernatant were subsequently transferred onto new, non-inflammatory MSC. After overnight culture, stromal cells were harvested and analyzed by RNA sequencing. Only when cultured with iMSC-conditioned neutrophils, stromal cells increased transcription of iMSC-defining genes such as neutrophil-recruiting chemokines *CXCL2*, *CXCL3* and *CXCL8*, plasma cell survival factors *IL6* and *LIF*, myeloid modulation factor *C3* and immune regulator *PTGS2* (Figures 4G-H, Supplemental Table 10), recapitulating the transcriptome of iMSCs in patients and induced iMSC-like cells *in vitro* (Figure 4I). Increased transcription of *CXCL8* and *IL6* in MSC cultured with iMSC-conditioned neutrophils was validated by qPCR (Supplemental Figure 4A). Levels of neutrophil-restricted transcripts were low and similar across conditions, excluding significant contamination of neutrophils in the stromal analysis (Supplemental Figure 4B). Moreover, induction of an iMSC-like transcriptome was independent from iMSC-secreted factors from the primary culture, as MSC cultured in the supernatant of iMSC alone did not induce transcription of inflammatory genes (Supplemental Figures 4C-D). The exception to this was *C3*, which is also associated with autocrine activation loops in fibroblasts from chronic inflammatory diseases^38^. As IL-1β is a potent inducer of iMSC, we tested whether MSC activation by neutrophils was IL-1β dependent by neutralizing IL-1β during co-culture. IL-1β neutralization reduced stromal activation by activated neutrophils, highlighting the importance of this pathway (Figures 4J-K, Supplemental Figure 4A, Supplemental Table 11). Combined, these data suggest that iMSC-activated neutrophils gain the ability to amplify stromal inflammation in an IL-1β-dependent manner.

### Residual stromal inflammation is present after first-line treatment

Despite intensive treatment regimens, virtually all myeloma patients experience disease relapse, even those in full remission after therapy. We hypothesized that persistence of pro-tumor inflammation could be implicated in recurrence of disease. Therefore, we set out to investigate presence of inflammatory stromal – neutrophil interactions in patients that completed first-line treatment. This included induction therapy, high-dose melphalan, autologous stem cell transplant (ASCT), and consolidation therapy with triplet alone or triplet + anti-CD38 monoclonal antibody therapy (Supplemental Table 1). First, we analyzed stromal cells in the post-treatment BM of MM patients. We previously identified the presence of stromal inflammation post-induction treatment^4^, and now set out to generate a single cell transcriptomic overview of the non-hematopoietic compartment of 5 patients after ASCT and consolidation therapy. Non-hematopoietic cells were isolated and sequenced as described before^4^, and subsequent integration with control data of non-hematopoietic cells previously generated in our lab^4^ created a dataset of 14,697 cells, of which 8,611 cells were derived from treated patients (Figure 5A). This dataset included five clusters of MSCs (MSC1-MSC5), a cluster of endothelial cells (EC), and a small cluster of *SELP*^+^ cells (SEC, Figure 5A-B, Supplemental Figure 4E). In contrast to NDMM, where transcription of inflammatory genes drove formation of separate clusters of iMSCs^4^, no specific inflammatory clusters were detected post-consolidation (Figure 5A). However, transcription of iMSC-defining genes remained elevated after consolidation treatment (Figure 5C, Supplemental Table 12). These genes included MM cell survival factors *IL6* and *LIF*, chemokines *CXCL2*, *CXCL3*, *CXCL5* and *CXCL8*, myeloid-modulating factors *C3*, *CCL2* and *ANXA1* and *CD44*^4, 39, 40^. Additionally, percentages of MSCs transcribing *CXCL2*, *CCL2*, *ANXA1* and *CD44* were significantly increased in BM aspirates of treated patients compared to controls (Figure 5D). The increased transcription of the immune modulator *PTGS2,* which defined iMSC at diagnosis, was lost after treatment (Figures 5D, Supplemental Figure 4F). Although no MM-specific clusters of iMSC were detected, transcription of inflammatory genes was mostly confined to cluster MSC2, which showed a trend towards enrichment in the treated BM compared to controls (Figure 5E, Supplemental Figures 4G-H). When translating NDMM-associated iMSC-specific gene transcription into an iMSC-like module score, this score was higher in MSCs of patients after consolidation treatment compared to controls (Figure 5F-G). Although the small size of our current cohort precludes definitive conclusions, no differences in fractions of iMSC-like cells were observed when comparing patients that underwent triplet-therapy with or without anti-CD38 antibodies (n = 2 vs. n = 3, Supplemental Figure 4I), or when comparing patients based on remission-status (n = 3 vs. n = 2, Supplemental Figure 4J). Finally, we used flow cytometric numeration of CD44-expressing MSCs as a proxy for the presence of iMSCs^4^. These experiments validated the presence of residual stromal inflammation in treated MM patients and confirmed the effect of treatment on reducing iMSC presence compared to NDMM (Figure 5H). In conclusion, first-line treatment reduces iMSC number and activation, but does not normalize stromal BM inflammation.

**Figure 5.**
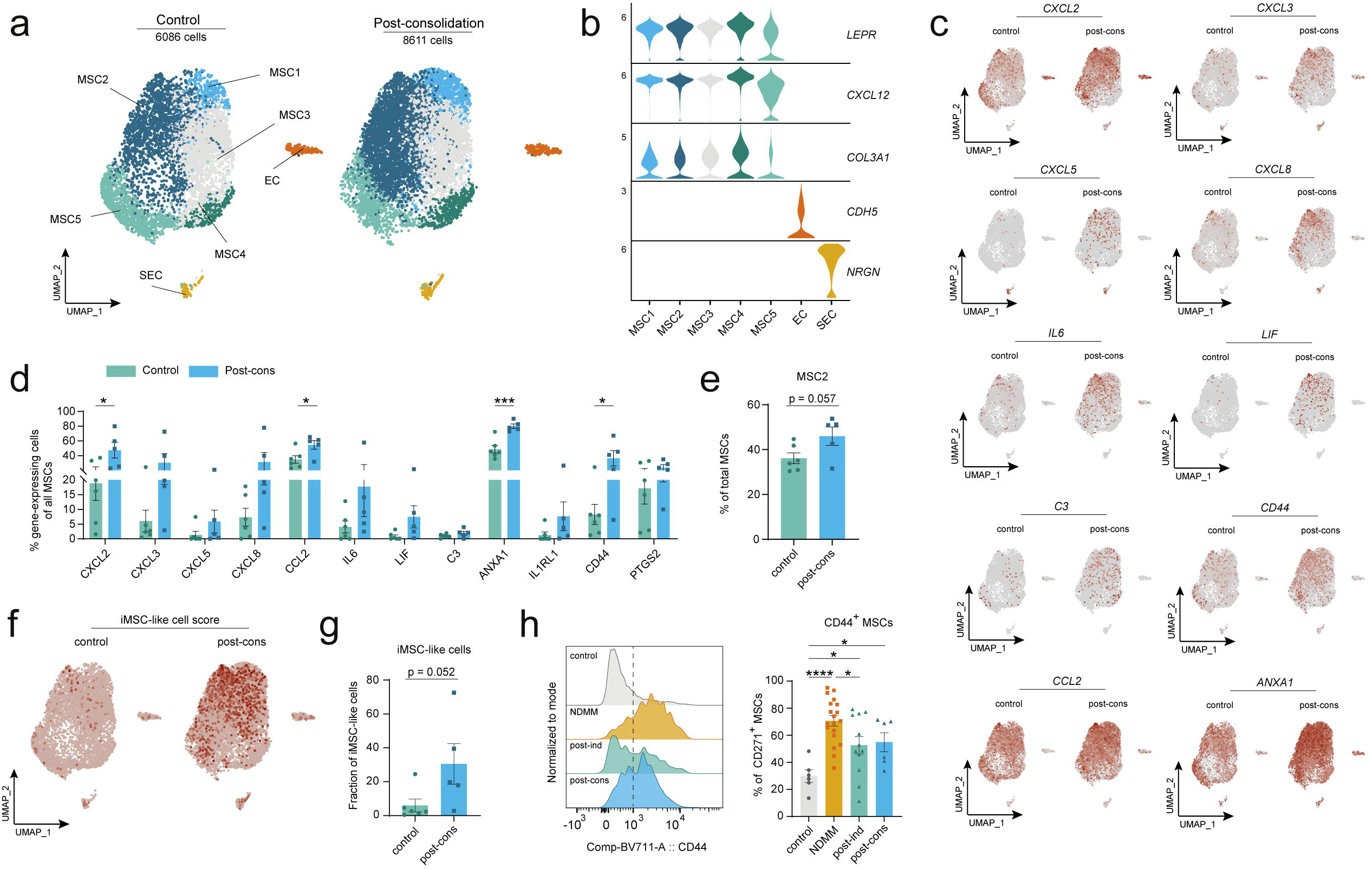
Residual stromal inflammation persists after first-line treatment. **A)** UMAP split in 8,611 cells from 5 treated MM patients and 6,066 cells from 6 non-cancer controls showing 7 clusters identified by markers genes in *(B)*. Colors represent clusters **B)** Marker gene transcription in the 7 clusters **C)** Transcription of selected differentially expressed genes in control or treated MM non-hematopoietic bone marrow cells **D)** Frequencies of MSC transcribing iMSC-associated genes per individual in each condition **E)** Fraction of MSC in cluster MSC2 per individual in each condition **F)** Expression of ‘iMSC-like cell’ score per cell in non-cancer controls and treated MM patients **G)** Frequencies of cells positive for ‘iMSC-like cell’ score per individual in each condition **H)** Representative histograms of CD44 expression in CD271^+^ MSCs of non-cancer controls, NDMM patients, and treated MM patients. Percentages of CD271^+^ cells positive for CD44 were quantified per individual in each condition. iMSC, inflammatory mesenchymal stromal cell. Significance was calculated in **D**, **E**, **G** and **H** using the Mann-Whitney *U* test (two-tailed); * P ≤ 0.05, ***P ≤ 0.001, ****P ≤ 0.0001.

### Pro-tumor neutrophils are reactivated after first-line treatment

Patients become neutropenic after high dose melphalan and ASCT^41^. However, the neutrophil compartment has recovered in numbers after consolidation treatment, as mature neutrophils were yet again the most prevalent nucleated cell in BM aspirates of post-treatment patients (Figure 6A). To assess the impact of anti-myeloma treatment on the pro-tumor phenotype of neutrophils, we isolated the myeloid compartment from BM aspirates of 5 patients that had completed first-line treatment (Supplemental Table 1). After single cell sequencing, the resulting dataset was integrated with the 4 non-cancer controls of Figure 1, and the neutrophilic lineage was isolated *in silico* (see methods). The complete neutrophil lineage was represented, with mature cells segregating in three clusters (MatNeu1, 2a and 2b; Figure 6B, Supplemental Figure 5A). Two clusters of mature neutrophils (MatNeu2a and MatNeu2b) were still characterized by transcription of pro-inflammatory and activation-related genes, including plasma cell survival factors, inflammasome components and ligands for inhibitory checkpoint molecules (Supplemental Figure 5B). Cluster MatNeu2b was distinguished from cluster MatNeu2a by presence of the IFN-response signature that was also seen in mature neutrophils at diagnosis (Figure 6C). Even though there was no longer an increase in percentage of cells in activated mature neutrophil clusters after treatment (Figures 6D-E), almost all pro-inflammatory genes were still increased in clusters MatNeu2a and MatNeu2b of MM patients compared to controls (Figure 6F, Supplemental Table 13). Importantly, mature neutrophils retained elevated levels of *TNFSF13B* and *IL1B* (Figures 6F-H, Supplemental Figure 5B) and, compared to other myeloid cell subsets post-consolidation, activated mature neutrophils remained prominent sources of these cytokines (Figure 6I). Similar to NDMM, highest transcription of *IL1B* was found in macrophages, yet the abundance of mature neutrophils (Figure 6A) suggests that macrophages and neutrophils both contribute to IL-1β production post-consolidation. Finally, mature neutrophils retain signs of STAT3 signaling after consolidation therapy, as clusters MatNeu2a/2b transcribed increased levels of *STAT3* and STAT3-target genes *SOCS3* and *BCL3* compared to non-cancer controls (Figure 6J). As most BM neutrophils post-consolidation will have matured after transplant, these data indicate that BM neutrophils are reactivated, potentially by residual iMSC, after first-line treatment.

**Figure 6.**
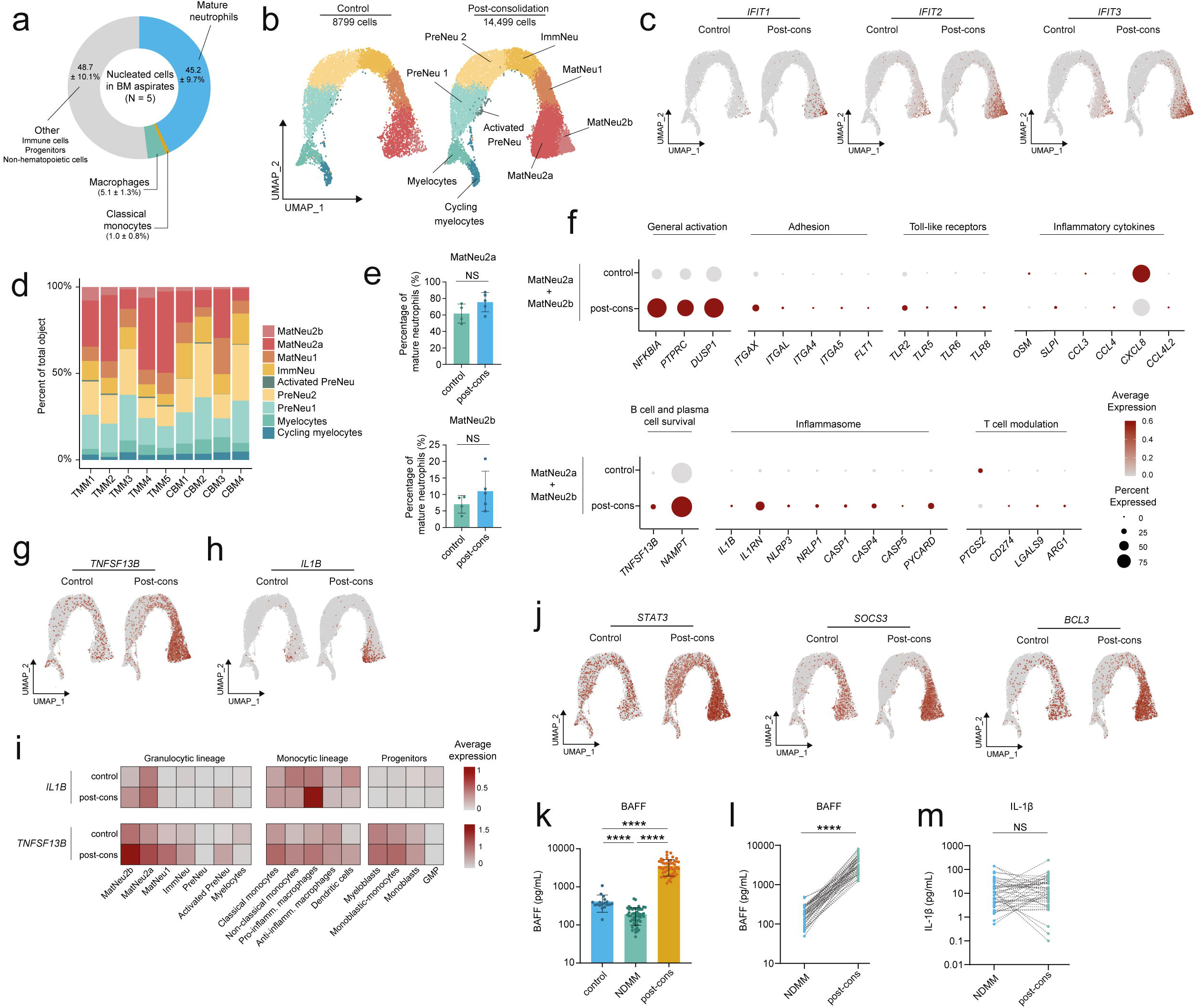
Pro-tumor neutrophils are reactivated after first-line treatment. **A)** Distribution of nucleated cells in BM aspirates of n = 5 MM patients after consolidation therapy, values are mean ± SD **B)** UMAP split in 14,499 cells from 5 MM patients after consolidation therapy and 8,799 cells from 4 non-cancer controls showing 9 clusters identified by marker genes in *Supplemental* Figure 3A. Colors represent clusters **C)** Transcription of interferon-response genes in non-cancer controls and treated MM patients (‘post-cons’) **D)** Cluster distribution per patient. Colors represent clusters. ‘TMM’, treated MM patient. ‘CBM’, control **E)** Percentages of mature neutrophils in clusters MatNeu2a and MatNeu2b per individual in each condition **F)** Transcription of differentially expressed genes in clusters MatNeu2a and MatNeu2b comparing controls to treated MM patients. Size of dot represents fraction of cells transcribing the gene, color represents level of transcription **G)** Transcription of *TNFSF13B* and **H)** *IL1B* in non-cancer controls and treated MM patients **I)** Average transcription of *IL1B* and *TNFSF13B* per annotated cluster of single cell RNA sequencing datasets. Data is split in average expression in non-cancer controls and treated MM patients **J)** Transcription of *STAT3*, *SOCS3* and *BCL3* in cells comparing control individuals and treated MM patients **K)** BAFF protein levels in BM plasma samples of N = 18 controls, and paired samples of N = 52 patients at diagnosis and after consolidation therapy **L)** BAFF protein levels in paired BM plasma samples of N = 52 MM patients at diagnosis and after consolidation therapy **M)** IL-1β protein in paired BM plasma samples of N = 52 MM patients. Post-cons, post-consolidation. Data are presented as mean ± SEM. Significance was calculated in **E** and **K** using the Mann–Whitney U test (two-tailed), and in **L** and **M** using the Wilcoxon Rank Sum test (two-tailed); ****P ≤ 0.0001, NS P > 0.05.

The increased transcription of *TNFSF13B* and *IL1B* in neutrophils after treatment may have consequences for disease recurrence. To assess if increased transcription also resulted in elevated protein production, we measured BAFF and IL-1β protein in paired BM plasma samples of 52 patients at diagnosis and after induction/consolidation treatment with triplet + anti-CD38 monoclonal antibody therapy (Supplemental Table 14). Surprisingly, BM plasma levels of BAFF were lower in MM patients at diagnosis than in non-cancer controls (190 ± 92 pg/mL vs. 411 ± 198 pg/mL, p <0.0001, N = 18 controls, Figure 6K, Supplemental Figure 5C). BAFF levels increased significantly after induction-therapy induced tumor debulking (Supplemental Figure 5C), raising the possibility that BAFF consumption by BCMA-expressing MM cells impacts measurable levels of BAFF at diagnosis. This notion is supported by significant reduction of BAFF levels in cultures of BAFF-producing neutrophils and MM cell lines (Supplemental Figure 3G). This notwithstanding, after consolidation treatment, when most of the tumor burden has been removed and BAFF consumption is no longer impacting BAFF measurements, levels of BAFF in BM were significantly elevated compared to non-cancer controls (3441 ± 1626 pg/mL vs. 411 ± 198 pg/mL, p <0.0001, Figure 6K-L). This increase in BAFF protein level was independent of remission status (n = 38 [73%] in full remission, Supplemental Figure 5D) or therapy response (n = 49 [92%] very good partial response [VGPR] or better, Supplemental Figure 5E). We did detect slightly lower levels of BAFF in patients that remained MRD-positive post-consolidation or had only partial treatment response (Supplemental Figures 5D-E), likely again reflecting BAFF consumption by residual tumor cells. We had previously shown that IL-1β is elevated in BM plasma of newly diagnosed patients^4^. After consolidation treatment IL-1β protein levels remained elevated (Figure 6M). This was most prominent in patients that remained MRD-positive or did not have a complete treatment response (Supplemental Figures 5F-G). In conclusion, reactivated neutrophils remain important sources of BAFF and IL-1β, reflected in elevated levels of these proteins in treated BM.

## Discussion

Neutrophils are increasingly acknowledged for their cancer-promoting abilities, and by virtue of their abundance in the BM, a pro-tumor bias of these cells could have a significant impact on BM-residing malignancies such as myeloma. Here, we provide evidence that in MM, inflamed stromal cells activate mature BM neutrophils towards a pro-inflammatory phenotype, including secretion of the BCMA-ligand BAFF and the pro-inflammatory cytokine IL-1β, factors with demonstrated adverse effects in MM^30, 31, 33, 34, 42^. Reciprocally, activated neutrophils can induce *de novo* stromal inflammation in an IL-1β dependent manner. This neutrophil-stromal cell axis persists after treatment, as both activated neutrophils and iMSC are present in patients that have undergone induction therapy, high-dose melphalan, ASCT and consolidation therapy (Supplemental Figure 6). Nevertheless, intensive treatment did have a noteworthy impact on BM inflammation in NDMM patients. Having previously established that iMSC remain present after induction treatment^4^, we now show that completion of initial treatment strongly reduced BM inflammation. However, even after this intensive treatment, and even in MRD-negative patients, the BM retains elevated numbers of iMSC compared to controls. It will be of interest to determine temporal evolution of BM inflammation in patients during maintenance treatment as soon as these samples become available.

Neutrophils are depleted by anti-MM therapy^41^, indicating that the pro-inflammatory neutrophils present after first-line therapy are the result of reactivation of newly formed cells. We entertain the hypothesis that persistence of residual iMSC after treatment leads to reactivation of newly-developing neutrophils post-ASCT. On the other hand, we cannot exclude that activation of newly-developing neutrophils is driven by therapy-related tissue damage. In such a scenario, neutrophil activation during consolidation treatment would maintain or re-inflame the stromal micro-environment, leading to persistence of BM inflammation. The ultimate conclusion of either scenario will be that both stromal and neutrophil activation need to be targeted in order to halt BM inflammation.

The TNF family member BAFF is a ligand for myeloma-cell expressed BCMA^25, 26^, a receptor involved in proliferation and survival of primary myeloma cells and myeloma cell lines *in vitro*^34, 43^. BAFF is a strong contributor to disease burden in myeloma mouse models^44^, and associated with rapid progression in patients^27^. Serum levels of BAFF are inversely correlated with MM patient survival^33^, signifying the importance of this growth factor in myeloma pathobiology. BAFF production in the MM BM has previously been ascribed to monocytes^45^, neutrophils^45^, myeloma cells^46^, osteoclasts^45^, and MSCs^43^. Here, using our in-house generated single cell sequencing datasets of the entire MM TME, we show that activated mature neutrophils transcribe the highest levels of *TNFSF13B*/BAFF. As they are also one of the most abundant cell types in the BM, mature neutrophils are likely a major source of BAFF in MM. Interestingly, BAFF transcription by activated neutrophils and BAFF BM plasma levels were still elevated after ASCT and consolidation therapy. In our current analyses, follow up was too short to associate presence of activated neutrophils and elevated BAFF levels with clinical outcomes. Analysis of neutrophil activation and concomitant increased BAFF-levels in larger patient cohorts and with prolonged follow-up will be an important step towards a full understanding of BM inflammation in MM.

Secretion of IL-1β in MM has previously been attributed to macrophages^31, 47^. However neutrophils also showed increased transcription of *IL1B* and elevated protein levels of intracellular (pro-)IL-1β, which makes it conceivable that neutrophils significantly contribute to the increased levels of IL-1β found in MM BM^4^. Interestingly, IL-1β is a main driver of IL-6 production in the MM tumor environment^48^, which is in line with the IL-1β-induced *IL6* transcription by iMSC shown in this study and previous work^4^. Carriers of polymorphic variants associated with reduced *IL1B* transcription are at lower risk for developing MM^49^ and a phase II trial investigating treatment of patients with smoldering multiple myeloma (SMM) or indolent MM with IL-1 receptor antagonist Anakinra demonstrated improved PFS^30^. Intriguingly, iMSC can induce IL-1β secretion by neutrophils, making this cytokine an attractive target to reduce BM inflammation, and by extension, provide a novel avenue for MM therapy development.

Stromal cells are known to impact tissue neutrophil function in cancer and during chronic inflammation^7, 50,51,52^. Here, we show that iMSC can induce an inflammatory profile in neutrophils that mimics BM neutrophils in MM patients. BM neutrophils from MM patients showed evidence of STAT3 activation and *in vitro*, neutrophil activation by inflammatory stromal cells was STAT3 dependent. Although our analysis did not lead to identification of the iMSC-derived signals activating neutrophils via STAT3, the STAT3 dependency may provide opportunity for therapeutic intervention of pro-tumor inflammation in the MM bone marrow. Multiple STAT3-inhibitors are currently under investigation for their application in cancer^53^, including myeloma^54^, where STAT3 inhibition additionally inhibits proliferation of myeloma cells^55, 56^.

In line with the previously described immune suppressive function of BM neutrophils in MM^14,15,16^, we show that mature neutrophils in the BM of NDMM and treated patients have increased transcription of genes associated with T cell inhibition. Interestingly, neutrophils did not acquire the T cell-inhibitory transcriptome when co-cultured with iMSC, while a previous study using MSC from patients with MM, or LPS-activated MSC from controls, demonstrated induction of neutrophils capable of functional T cell inhibition^19^. This discrepancy could be a limitation of our transcriptional analysis, or might be driven by signals beyond our current analysis.

In summary, our work identifies neutrophils as core components of BM inflammation by providing the MM TME with survival factor BAFF and disease-promoting IL-1β. We conceptualize that neutrophil activation is mediated by MM-specific stromal inflammation, creating a self-perpetuating inflammatory loop that persists even after first-line treatment and merits future investigation into the role of this stromal cell-neutrophil axis in disease recurrence.

## Author contributions

Conceptualization, M.D.J and T.C.; Methodology, M.D.J. and T.C.; Investigation, M.D.J., C.F, N.P., M.V., T.V.H., S.T., and M.B.H.; Formal analysis, M.D.J., C.F., N.P., M.V., T.V.H., R.H., G.V.B., M.A.S., M.B.H., H.B., and T.C.; Resources, P.V.D.W, P.M., F.G., M.V.D., A.B. and P.S.; Data curation, M.D.J., R.H. and G.V.B. ; Visualization, M.D.J.; Writing – Original Draft, M.D.J. and T.C.; Writing – Review & Editing, all authors; Funding Acquisition, T.C. and P.S.; Supervision, T.C., and P.S.; Project administration, T.C, and P.S.

## Materials and methods

### Sample acquisition

For studies of the myeloid compartment, we obtained fresh BM aspirates at diagnosis and after consolidation treatment from myeloma patients included in trials sponsored by the European Myeloma Network (EMN). For studies of the non-hematopoietic compartment, frozen BM aspirates were obtained from myeloma patients after consolidation treatment from the Cassiopeia study (Intergroupe Francophone du Myeloma (IFM) 2015-01/Dutch-Belgian Cooperative Trial Group for Hemato-Oncology [HOVON]131; ClinicalTrials.gov Identifier: NCT02541383). Marrow aspirates were used within 24 hours after collection. Paired bone marrow plasma was also obtained from the Cassiopeia study. Naive neutrophils were collected from peripheral blood of healthy individuals. All material was obtained after written informed consent approved by the Institutional Review Boards of the Erasmus Medical Center (Rotterdam, the Netherlands) in accordance with the Declaration of Helsinki.

### Cell lines

Human adipose-derived stromal cells (ADSCs) were acquired from Thermo Fisher, cultured in complete MesenPRO RS medium (Gibco) supplemented with 1% penicillin/streptomycin, and used until passage 9. During experiments, ADSCs were cultured in DMEM supplemented with 10% fetal calf serum (FCS, Corning) and 1% penicillin/streptomycin (Sigma-Aldrich). MM1S and U266 cells were acquired from ATCC, and cultured in RPMI 1640 supplemented with 10% FCS (Corning), 200nM L-glutamine (Gibco) and 1% penicillin/streptomycin (Sigma-Aldrich).

### Isolation of BM myeloid cells for transcriptomic analyses

To remove erythrocytes, unfractionated BM samples were incubated with red blood cell (RBC) lysis buffer (0.15M NH_4_Cl, 10mM KHCO_3_, 0.1mM EDTA in miliQ water) at a ratio of 1:4 for 5min at room temperature. Fc receptors were blocked with 10% normal human AB serum (Sigma-Aldrich). After blocking, cells were stained for sorting in PBS containing 2% FCS at 4°C with the following antibodies: CD138-FITC (1:20; MI15, BioLegend), CD45-PerCP-Cy5.5 (1:20; 2D1, eBioscience), CD11b-PE (1:20; ICRF44, eBioscience), CD14-PECy7 (1:20; HCD14, BioLegend), CD163-PE-Dazzle594 (1:50; GHI/61, BioLegend), CD3-APCCy7 (1:20; HIT3a, BioLegend), CD71-Alexa Fluor 700 (1:20; Mem-75, ExBio), CD235a-APC (1:100; GA-R2, BD Biosciences), CD16-BV510 (1:70; 3G8, BioLegend). DAPI (Life Technologies) was used for dead cell exclusion. Granulocytes were sorted as CD45^mid/hi^CD138^-^CD235a^—^CD71^—^ CD3^—^ cells within the granulocytic scatter. Mononuclear phagocytes were sorted as CD45^mid/hi^CD138^—^CD235a^—^CD71^mid/—^CD3^—^ cells within the monocytic scatter. Cells were sorted in DMEM + 10% FCS using a FACSAria III (BD Biosciences) and BD FACSDiva version 5.0 (BD Biosciences).

### Isolation of non-hematopoietic cells for transcriptomic analyses

Non-hematopoietic cells were isolated as described before^4^. Mononuclear cell fractions were isolated using Histopaque (Sigma-Aldrich), viably frozen in Dulbecco’s modified eagle medium (DMEM, Gibco) supplemented with 40% heat-inactivated fetal calf serum (FCS, Corning) and 10% dimethyl sulfoxide (DMSO, Sigma-Aldrich), and stored in liquid nitrogen until use. Viably frozen bone marrow aspirates were thawed in DMEM supplemented with 10% FCS according to the 10x Genomics protocol ‘Fresh Frozen Human Peripheral Blood Mononuclear Cells for Single-cell RNA sequencing’. Samples containing >150 × 10^6^ mononuclear cells were used for non-hematopoietic cell isolation. Fc receptors were blocked with 10% normal human AB serum (Sigma-Aldrich). To enrich for MSCs, samples were incubated with biotinylated antibodies against CD45 (1:50, clone HI30, BioLegend), CD235a (1:50, clone HIR2, BioLegend) and CD38 (20 µl per 10^7^ cells, from human CD38 MicroBead kit, Miltenyi Biotec) followed by depletion using magnetic anti-biotin beads (30 µl per 10^7^ cells, Miltenyi Biotec) and the iMag (BD Biosciences). After depletion, cells were stained for sorting in PBS containing 2% FCS at 4 °C with the following antibodies: CD271-PE (1:200; ME20.4, BioLegend), CD235a-PECy7 (1:20; HI264, BioLegend), CD31-APCCy7 (1:20; WM59, BioLegend), CD44-BV711 (1:50; IM7, BioLegend), streptavidin-BV421 (1:200; BioLegend), CD34-PECF610 (1:100; 4H11, eBioscience), CD45-APC (1:20; 2D1, eBioscience), CD38-FITC (1:20; MHCD3801, Life Technologies), CD71-Alexa Fluor 700 (1:20; MEM-75, ExBio) and CD105-BV510 (1:50; 266, BD Biosciences). 7AAD (Beckman Coulter) was used for dead cell exclusion. For single-cell sequencing studies, CD45^—^CD38^—^CD235a^—^CD71^—^ non-hematopoietic cells were sorted in DMEM supplemented with 10% FCS using a FACSAria III (BD Biosciences) and BD FACSDiva version 5.0 (BD Biosciences).

### Single cell RNA sequencing

Single cells were encapsulated for cDNA synthesis and barcoded using the Chromium Single Cell 3’ Reagent Kit v3.1 (10X Genomics), followed by library construction according to the manufacturer’s recommendations. For the generation of the myeloid dataset, sorted granulocytes were combined with sorted mononuclear phagocytes at a 1:1 ratio and processed as one sample. For the generation of the non-hematopoietic dataset, sorted non-hematopoietic cells were combined with CD45^+^CD38^—^CD34^—^ sorted hematopoietic cells and processed as one sample. The quality and quantity of libraries was determined using the Agilent 2100 Bio-Analyzer and the High Sensitivity DNA kit. Libraries were sequenced on the NovaSeq 6000 platform (Illumina), paired-end mode, at a sequencing depth of ∼20.000 reads/cell, followed by computational alignment using CellRanger (v 6.0.1, 10x Genomics).

### Single cell transcriptome analyses

Datasets were subjected to quality control steps using Seurat (v 4.0.1)^57^ that included selecting cells with a library complexity of more than 100 features, removing doublets (cells with double amounts of UMIs), and filtering out cells with high percentages of mitochondrial genes (>10%)(see https://github.com/MyelomaRotterdam for specifics).

For the myeloid datasets, data from samples of 6 (NDMM) or 5 (post-consolidation) patients and 4 non-cancer controls were normalized, merged by integration and label transfer^58^ and analyzed by principal-component analysis (PCA) on the most variable genes (k = 2,000) across all cells. With the principal components (PCs) (k = 10-20), unsupervised clustering was performed using a shared nearest neighbor (SNN) modularity optimization-based clustering algorithm (resolution 0.3)^59^ and cells were projected in two dimensions using Uniform Manifold Approximation and Projection (UMAP)^60^. Pseudotime analysis was carried out using Monocle 3^61^. Progenitors, mononuclear phagocytes and neutrophilic lineages were separated into three datasets by retrieving replicate-specific barcodes and rerunning the integration (see https://github.com/MyelomaRotterdam for specifics). Cluster annotation was as follows: GMP, *CD34*^+^*MKI67*^+^*MPO*^—^, myeloblasts, *MPO*^+^*ELANE*^+^*CD34*^—^; monoblasts, *MPO*^+^*CD14*^+^*CD34*^—^; cycling progenitors, *MKI67*^+^*MPO*^+^*CD34*^—^; classical monocytes, *CD14*^+^*FCGR3A*^—^; non-classical monocytes, *CD14*^—^*FCGR3A*^+^; pro-inflammatory macrophages, *CD163*^+^*IL1B*^+^; anti-inflammatory macrophages, *CD163*^+^*LTF*^+^; dendritic cells, *CD1C*^+^; myelocytes, *CD14*^—^ *ELANE*^+^*AZU1*^+^; pre-neutrophils *CD14*^—^*MPO*^—^*LTF*^+^; immature neutrophils *ARG1*^+^*MMP9*^+^; mature neutrophils, *MME*^+^*FCGR3A*^+^.

For the non-hematopoietic dataset, data from samples of 5 patients after consolidation therapy was integrated with data of 6 (of a total of 7) non-cancer controls cells generated previously^4^. The 7^th^ control (‘CBM7’ in ^4^) had generalized periodontitis and paradontitis at the time of sampling, which can affect the immune system systemically^62^, and was therefore excluded. After normalization and principal-component analysis (k = 10-20) on the most variable genes (k = 2,000), cells expressing *CXCL12* (MSCs), *LEPR* (MSCs), *CDH5* (endothelial cells), *RUNX2* (osteolineage) or *SELP* (*SELP*^+^ cells) were separated *in silico* by retrieving replicate-specific barcodes, which were used to rerun the integration (see https://github.com/MyelomaRotterdam for specifics). With the principal components (PCs) (k = 10), unsupervised clustering was performed using a shared nearest neighbor (SNN) modularity optimization-based clustering algorithm (resolution 0.3)^59^ and cells were projected in two dimensions using UMAP. The iMSC-like cell score was computed using Seurat’s AddModuleScore() function using the top 50 DEGs of iMSC detected in NDMM (positive only)^4^. Cluster annotation was as follows: mesenchymal stromal cells (MSC), *LEPR*^+^*CXCL12*^+^*COL3A1*^+^; endothelial cells (EC), *CDH5*^+^*CD36*^+^; P-selectin^+^ cells (SELP), *SELP*^+^*NRGN*^+^*CXCL12*^—^*CDH5*^—^.

To analyze bone marrow-wide transcription of *TNFSF13B*/*IL1B*, all in-house datasets were combined by integration and label-transfer. Although new clusters were formed after scaling, principal-component analysis (k = 18) and unsupervised clustering (resolution = 0.5), subsequent analyses were carried out using the annotation of the original objects, prior to integration. Average expression of *TNFSF13B* and *IL1B* were calculated per cluster using Seurat’s AverageExpression() function. Heatmaps were generated using the pheatmap^63^ package.

Differentially expressed genes were determined with the Seurat FindMarkers function. Gene set enrichments were calculated with GSEA software (v 3.0, Broad Institute) using the pre-defined gene sets from the Molecular Signatures Database (MSigDB 6.2, H: hallmark gene sets)^64, 65^ or newly-constructed gene sets based on previously generated datasets. The gene list was ranked according to log2FCs from the FindMarkers function and the pre-ranked GSEA was run using the classic enrichment statistic with 1000 permutations. Frequency bar plots were generated using the DittoSeq package (v 1.10.0)^66^.

### Proliferation cultures

0.5 x 10^5^ U266 cells were seeded in round-bottom 96-well plates in RPMI, supplemented with 10% FCS, 200nM L-glutamine and 1% penicillin/streptomycin, with or without 200ng/mL recombinant human BAFF (R&D Systems). After overnight culture, cells were counted using the Countess™ 3 (Thermo Fisher).

### Immunofluorescence

Formalin-fixed, paraffin-embedded (FFPE) bone marrow specimens were retrieved from the archive of the Institute of Pathology, FAU Erlangen-Nuremberg, including 10 bone biopsies with plasma cell myeloma and 7 biopsies from patients with iron deficiency anemia as non-cancer controls. For antigen retrieval, slides were cooked in a steam cooker (Biocarta) for 2.5 minutes in TRS6 (Dako Cytomation). Then, slides were incubated overnight with anti-human IL-1β (1:100, 615417, R&D Systems) at room temperature, followed by development using the Polymer-Kit (Zytomed POLAP-100) and FastRed (Sigma-Aldrich). Thereafter, slides were incubated with anti-human CD15 (1:10, C3D-1, Diagnostic BioSystems) for 1 hour at 37°C. Fluorescence was developed by incubation with a secondary biotinylated goat-anti-mouse IgM antibody (1:500, M0806, Vector) for 1 hour at room temperature followed by incubation with streptavidin Alexa Fluor 488 (Life Technologies) for 1 hour at room temperature.

### Confocal microscopy

Imaging was performed using the LSM 700 confocal microscope (Zeiss) at a magnification of 630x. Regions of Interest (ROIs) matching granulocytes (CD15^+^) were segmented using Zeiss-software (ZEN 2.6), and the mean fluorescence intensity (MFI) of IL-1β was assessed for each ROI. Data from 25 granulocytes from each donor were used to determine frequencies of IL-1β positive cells.

### Protein analyses

Bone marrow supernatant was collected from bone marrow aspirates after Histopaque (Sigma-Aldrich) separation and stored at -80° until use. Prior to protein analysis analyses, supernatants vials were spun down twice at 21,000 x g for 5 min to pellet and discard lipid contents, debris and residual cells. Levels of BAFF protein were determined in culture supernatant or bone marrow plasma using the human BAFF/BLyS/TNFSF13B DuoSet ELISA kit (R&D Systems) in combination with the Pierce Streptavidin-Poly-HRP (1:10,000; Thermo Scientific). Levels of IL1β protein were determined in culture supernatant using the IL-1 beta/IL-1F2 DuoSet ELISA kit (R&D Systems) in combination with the Pierce Streptavidin-Poly-HRP (1:10,000; Thermo Scientific) and in bone marrow plasma using the Human IL-1 beta/IL-1F2 Quantikine HS ELISA Kit (R&D Systems).

### Naïve neutrophil isolation

Neutrophils were isolated from peripheral blood samples using the EasySep Direct Neutrophil Isolation kit (StemCell) in combination with the EasySep magnet, according to the manufacturer’s recommendations. Purity of neutrophils after isolation ranged from 96-98% based on location in FSC/SSC scatter. Enriched neutrophils were cultured in Iscove’s Modified Dulbecco Medium (IMDM) supplemented with 10% FCS, 500nM L-glutamine (Gibco), 50uM 2-mercaptoethanol (Sigma-Aldrich) and 1% penicillin/streptomycin (Sigma-Aldrich).

### Stromal-neutrophil co-cultures

iMSC-like cells were acquired by stimulating ADSCs with recombinant human IL-1β (0.5ng/ml, Miltenyi Biotec) overnight. Unstimulated ADSCs (‘MSCs’) were used as controls. The supernatant containing recombinant IL-1β was removed prior to the start of co-cultures. Coculture experiments were performed by adding 300,000 naïve neutrophils to 50-60% confluent MSC or iMSC-like cell monolayers in 96-well flat-bottom plates with or without addition of IL-1β (0.5ng/mL Miltenyi), G-CSF (2000IU/mL, filgrastim, Sandoz), LPS (0.5ug/mL, Sigma-Aldrich), Stattic (3.5uM, Abcam) or DMSO (WAK-Chemie Medical GmbH). After 24hrs, supernatants were collected for ELISA, and neutrophils were harvested by gentle pipetting to asses activation status and phosphorylation of STAT3 or lysed in lysis buffer containing TCEP (Macherey-Nagel) for RNA sequencing. To analyze the capacity of inflammatory neutrophils to activate MSC, conditioned neutrophils were harvested and pooled. 600,000 cells plus culture medium were replaced in a 24 well plate containing a 50-60% confluent monolayer of unstimulated ADSCs with or without addition of anti-human IL-1β (2ug/mL, R&D Systems) or a mouse IgG1 isotype control (2ug/mL, R&D Systems). After overnight culture, neutrophils were removed by gentle pipetting, and ADSCs were lysed in the plate using in lysis buffer containing TCEP (Macherey-Nagel). Lysed cells were snap frozen in nitrogen and stored at -80°C until use.

### Flow cytometry

To asses activation status of neutrophils after co-culture, harvested neutrophils were resupended in PBS containing 2% FCS and Fc receptors were blocked with 10% normal human AB serum (Sigma-Aldrich). Cells were stained with the following antibodies: CD66b-FITC (1:250; G10F5, BioLegend), CXCR1-PerCP (1:50; 8F1/CXCR1, BioLegend), CD11b(act)-PECy7 (1:100; CBRM1/5, BioLegend), CD11c-APCCy7 (1:100; Bu15, BioLegend), CD45-Alexa Fluor 700 (1:100; HI30, BioLegend), C3AR-APC (1:10; hC3aRZ8, BioLegend), CD10-BV510 (1:50; HI10a, BioLegend) and CD11b-BV711 (1:50; ICRF44, BioLegend). DAPI (Life Technologies) was used for dead cell exclusion. To asses phosphorylation of STAT3, harvested neutrophils were resupended in PBS + 2% FCS and Fc receptors were blocked with 10% normal human AB serum (Sigma-Aldrich). Cells were stained with CD15-APCCy7 (1:100; W6D3, BioLegend) and LIVE/DEAD Fixable Violet stain (Invitrogen), and subsequently fixed in Cytofix fixation buffer (BD Biosciences) and washed twice. Permeabilisation was carried out at -20°C in True-Phos Perm buffer (BioLegend), which was added dropwise to cells that were being vortexed. After permeabilisation, cells were washed twice and stained with pSTAT3[Tyr705]-PE (1:10; 13A3-1, BioLegend) at room temperature. Samples were measured on FACSymphony A5 (BD Biosciences). Analyses were carried out using FlowJo version 10.6.1 (BD Biosciences).

To asses surface molecule expression of BM neutrophils, unfractionated fresh BM samples were incubated with red blood cell (RBC) lysis buffer (0.15M NH4Cl, 10mM KHCO3, 0.1mM EDTA in miliQ water) at a ratio of 1:4 for 5min at room temperature. Fc receptors were blocked with 10% normal human AB serum (Sigma-Aldrich). For analysis of neutrophil activation, cells were stained in PBS containing 2% FCS at 4°C with the following antibodies: CXCR1-PerCP (1:50; 8F1/CXCR1, BioLegend), CD11b-PE (1:100; ICRF44, eBioscience), CD11b(act)-PECy7 (1:100; CBRM1/5, BioLegend), CD62L-PE Dazzle 594 (1:100; DREG-56, BioLegend), CD45-Alexa Fluor 700 (1:100; HI30, BioLegend) and CD10-BV510 (1:50; HI10a, BioLegend). For quantification of BM aspirate cell distribution and CXCR1/CXCR2 expression, cells were stained in PBS containing 2% FCS at 4°C with the following antibodies: CXCR1-FITC (1:100; 8F1/CXCR1, BioLegend), CD19-PerCP (1:50; HIB19, BioLegend), CD56-PE (1:20; MY31, BD Biosciences), CD14-PECy7 (1:20; HCD14, BioLegend), CXCR2-PE Dazzle 594 (1:50; 5E8/CXCR2, BioLegend), CD3-APCCy7 (1:20; HIT3a, BioLegend), CD34-Alexa Fluor 700 (1:20; 581, BioLegend), CD15-APC (1:50; HI98, BioLegend) and CD16-BV510 (1:300; 3G8, BioLegend). DAPI (Life Technologies) was used for dead cell exclusion. Samples were measured on a LSRII (BD Biosciences). Analyses were carried out using FlowJo version 10.6.1 (BD Biosciences).

For quantitative analysis of CD44^+^ iMSCs, cells were stained in PBS containing 2% FCS at 4°C with the following antibodies: CD235a-PECy7 (1:20; HI264, BioLegend), CD31-APCCy7 (1:20; WM59, BioLegend), CD44-BV711 (1:50; IM7, BioLegend), streptavidin-BV421 (1:200; BioLegend), CD34-PECF610 (1:100; 4H11, eBioscience), CD45-APC (1:20; 2D1, eBioscience), CD38-FITC (1:20; MHCD3801, Life Technologies), CD71-Alexa Fluor 700 (1:20; MEM-75, ExBio) and CD105-BV510 (1:50; 266, BD Biosciences). DAPI (Life Technologies) was used for dead cell exclusion. Samples were measured on a FACSAria III (BD BioSciences). Analyses were carried out using FlowJo version 10.6.1 (BD Biosciences).

### Bulk RNA sequencing

RNA was extracted from lysed cells using the NucleoSpin RNA XS kit (Machery-Nagel). cDNA was prepared using the SMARTer Ultra Low RNA kit for Illumina Sequencing (Clontech Laboratories) according to the manufacturer’s protocol. The quality and quantity of cDNA samples was verified using the Agilent 2100 Bio-Analyzer and the High Sensitivity DNA kit. cDNA Libraries were generated using the TruSeq Sample Preparation v.2 Guide (Illumina) and paired end-sequenced on the NovaSeq 6000 (Illumina, read length 2 x 101bp). Adaptor sequences and polyT tails were trimmed from unprocessed reads with fqtrim (v 0.9.7, http://ccb.jhu.edu/software/fqtrim/). Per transcript, pseudocounts and TPMs were determined with Salmon (v 1.4.0) ^67^ using the Hg38 genome reference (release 103). Subsequently, gene pseudocount estimates were determined using tximport (v 1.22.0)^68^. These per gene pseudocount estimates were in turn used for determining differential expression of genes between groups of interest through the DESeq2 (v 1.34.0) package^69^ based on Benjamini-Hochberg corrected p-values. GSEA was performed with the GSEA software (v 3.0, Broad Institute) using predefined gene sets from the Molecular Signatures Database (MSigDB 6.2, H: hallmark gene set)^65, 70^ or self-made gene sets from previously generated datasets. Gene lists were ranked on the basis of the log2fold-change made available through the DESeq2 package. Classical enrichment statistics with 1000 permutations were used to determine significant enrichment within gene sets. Heatmaps were generated using the pheatmap package (v 1.0.12)^63^, and volcano plots were generated using the EnhancedVolcano package (v 1.16.0)^71^.

### Transcript analyses

RNA was extracted using the NucleoSpin RNA XS kit (Machery-Nagel) and cDNA was generated using the SensiFast cDNA synthesis kit (BioLine). Quantitative PCR (qPCR) was carried out using a Neviti Thermal Cycler (Applied Biosystems) and DyNAmo Flash SYBR Green qPCR kit (Finnzymes), with the addition of MgCl_2_ to a final concentration of 4 mM. All reactions were performed in duplicate and were normalized to the expression of *GAPDH*. Relative expression was calculated by the cycling threshold (CT) method as -2^ΔCT^. Primer sequences were as follows: *GAPDH* (forward, 5′-GTCGGAGTCAACGGATT-3′; reverse, 5′-AAGCTTCCCGTTCTCAG-3′), *SOCS3* (forward, 5’-GCGCCTCAAGACCTTC-3’; reverse, 5’-CCCCCTCACACTGGAT-3’).

### Statistical analyses

Statistical analysis was carried out in R (v 4.1.0) or Prism 8 (GraphPad Software). Visualization was carried out using ggplot2 (v 3.4.1)^72^ or Prism 8 (GraphPad Software). All results in graphs are means ± SEM. Tests used to evaluate statistical significance are detailed in figure legends.

### Data availability

The single cell RNA-sequencing datasets described in this study are available on ArrayExpress no. E-MTAB-12760. CellRanger output files are available through Mendeley Data (https://data.mendeley.com/datasets/sm7fvt8hsg). Datasets can also be interactively explored at www.bmbrowser.org. The code generated during this study to analyze the single cell datasets is available through GitHub: https://github.com/MyelomaRotterdam. Bulk RNA-sequencing data of neutrophils and ADSCs acquired after co-culture are available on EGA no. EGAD00001010080. Further information and requests for resources and reagents should be directed to and will be fulfilled by the lead contact, Tom Cupedo.

## Acknowledgements

We thank members of Myeloma Research Rotterdam and the department of Hematology for critical discussions and reading of the manuscript, and patients, families, and nurses for their contributions to this study. We thank professor Ton Langerak (dept. of Immunology, Erasmus MC, Rotterdam) for provision of peripheral blood samples of normal controls. H.B. was supported by the Wilhelm-Sander Foundation. This work was supported by an International Myeloma Society and Paula and Rodger Riney foundation translational research award (T.C.), and grants of the European Myeloma Network (P.S.).

