## Supplemental Information for "An IL-1β driven neutrophil-stromal cell axis fosters a BAFF-rich microenvironment in multiple myeloma"

### Supplemental Figure 1

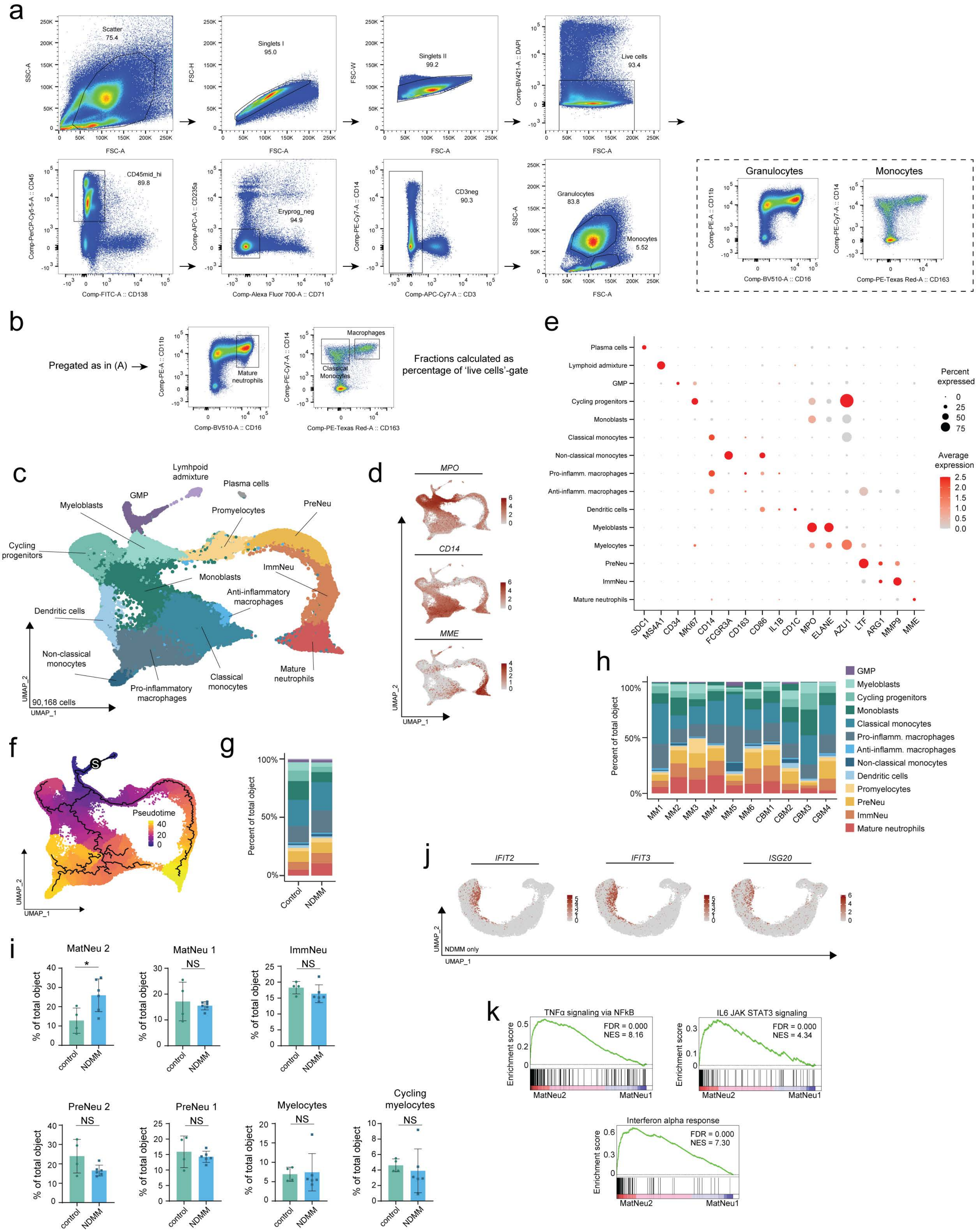

**Supplemental Figure 1** *Related to Figure 1* **A)** Gating strategy for neutrophils and monocytes/macrophages of unfractionated bone marrow from aspirates for single-cell RNA sequencing experiments **B)** Gating strategy for quantification of mature neutrophils, monocytes and macrophages in unfractionated bone marrow aspirates. Fractions were calculated as percentages of 'live cells'-gate (as depicted in *(A)*) **C)** UMAP of the combined dataset of 52,542 cells from NDMM patients and 37,626 cells from controls showing 15 clusters identified by marker genes in *(D)* and *(E)*. Colors represent clusters **D)** Marker gene transcription to demarcate the 3 major myeloid populations: progenitors (*MPO*$^{+}$), mononuclear phagocytes (*CD14*$^{+}$) and neutrophils (*MME*$^{+}$ marks mature neutrophils) **E)** Transcription of marker genes of the myeloid dataset **F)** Monocle 3 pseudotime analysis of the myeloid dataset. 'S' represents an arbitrarily selected starting point of trajectory calculations **G)** Cluster distribution per condition. Colors represent clusters **H)** Cluster distribution per patient. Colors represent clusters. 'MM', MM-patient. 'CBM', control **I)** Percentages of neutrophils in the 7 clusters per individual in each condition **J)** Transcription of interferon-response genes comparing non-cancer controls to NDMM patients **K)** Enrichment plots for selected Hallmark genesets comparing cluster MatNeu2 to cluster MatNeu1 in NDMM patients;
FSC, forward scatter. SSC, sideward scatter. GMP, granulocyte-monocyte progenitor. Data are presented as mean ± SEM. Significance was calculated in **I** using the Mann–Whitney U test (two-tailed), *P ≤ 0.05, NS P > 0.05.

### Supplemental Figure 2

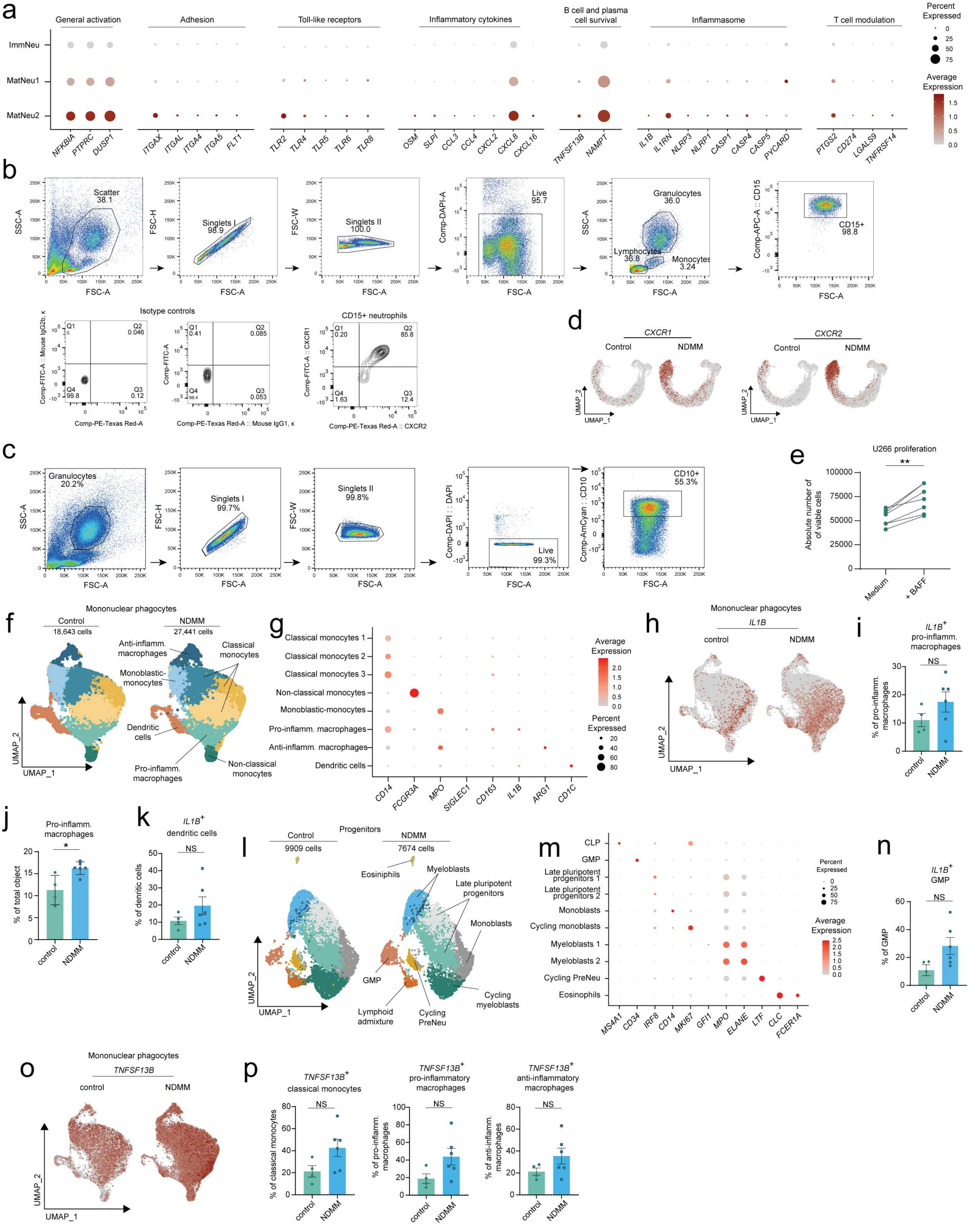

**Supplemental Figure 2** *Related to Figures 1 and 2* **A)** Transcription of selected genes comparing cluster MatNeu1 to clusters MatNeu2 and ImmNeu in NDMM patients **B)** Gating strategy for analysis of CXCR1/CXCR2 expression on (im)mature neutrophils, including FITC/PE fluorescence levels when samples were stained with relevant isotype controls and a representative plot of CXCR1/CXCR2 expression on $CD15^+$ neutrophils **C)** Pre-gating strategy for live $CD10^+$ neutrophils used for quantification of CD62L and CD11b(act) expression **D)** Transcription of *CXCR1* and *CXCR2* in controls and NDMM patients **E)** Absolute number of U266 myeloma cells cultured overnight in medium alone or medium supplemented with BAFF. Lines depict paired samples **F)** UMAP of the combined dataset of 27,441 mononuclear phagocytes from NDMM patients and 18,643 mononuclear phagocytes from control individuals showing 8 clusters identified by marker genes in *(G)*. Colors represent clusters **G)** Transcription of marker genes in the mononuclear phagocytes dataset **H)** Transcription of *IL1B* in mononuclear phagocytes of controls and NDMM patients **I)** Quantification of *IL1B*-transcribing cells in the cluster of pro-inflammatory macrophages per individual per condition **J)** Quantification of the percentage of cells in the total object that are in the cluster pro-inflammatory macrophages **K)** Quantification of *IL1B*-transcribing cells in the cluster of dendritic cells per individual per condition **L)** UMAP of the combined dataset of 7674 progenitors from NDMM patients and 9909 progenitors from control individuals showing 9 clusters identified by marker genes in *(M)*. Colors represent clusters **M)** Transcription of marker genes in the progenitor dataset **N)** Quantification of *IL1B*-transcribing cells in the cluster of GMP per individual per condition **O)** Transcription of *TNFSF13B* in mononuclear phagocytes of control individuals and NDMM patients **P)** Quantification of *TNFSF13B*-transcribing cells in the clusters of classical monocytes, pro-inflammatory macrophages and anti-inflammatory macrophages per individual per condition.

MHC, major histocompatibility complex. Data are presented as mean ± SEM. Significance was calculated in **I**, **J**, **K**, **N** and **P** using the Mann–Whitney U test (two-tailed), and in E using the Willcoxon Rank Sum test (two-tailed); *P ≤ 0.05, **P ≤ 0.01, NS P > 0.05.

### Supplemental Figure 3

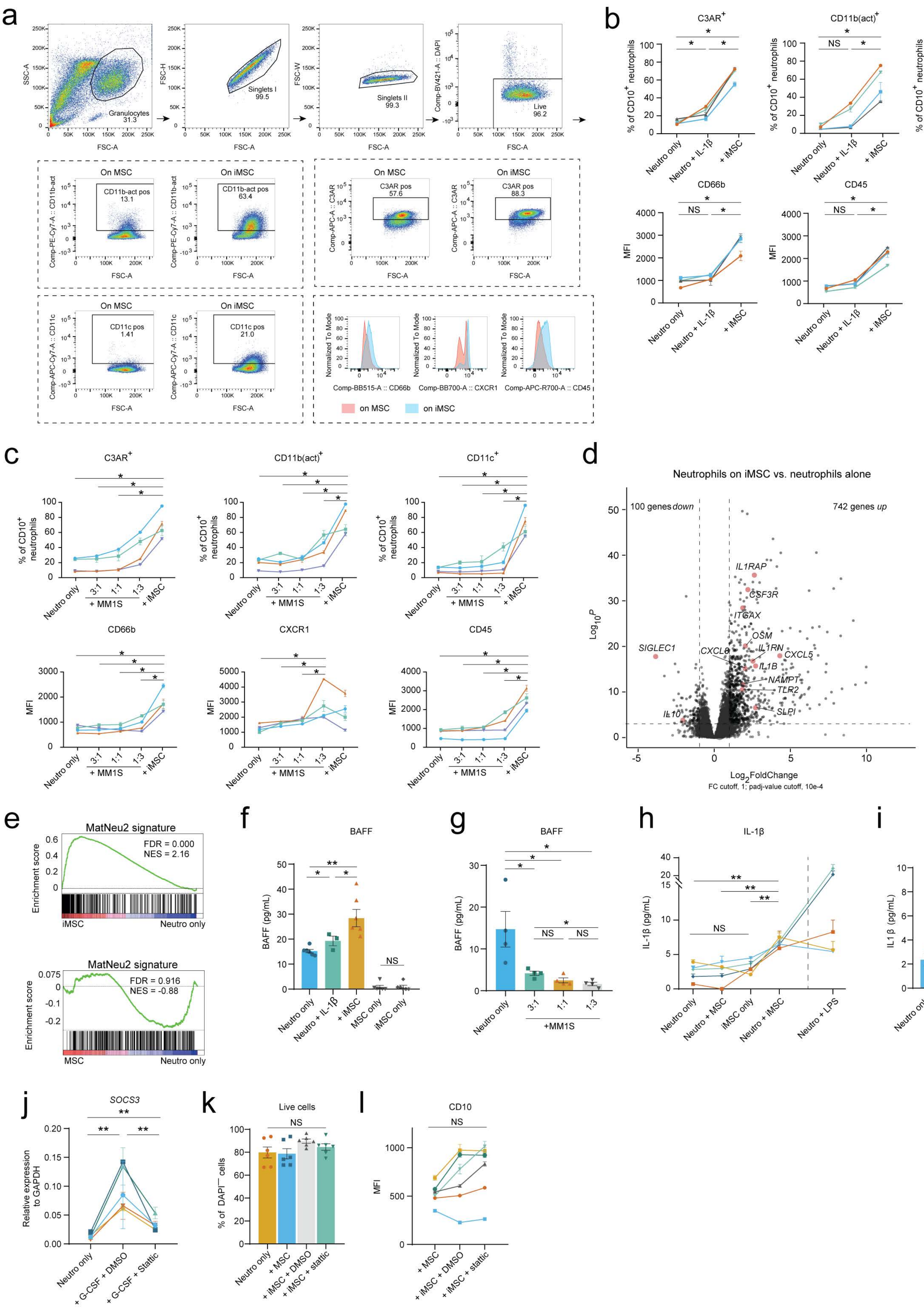

**Supplemental Figure 3** *Related to Figures 3 and 4* **A)** Gating strategy for analysis of activation marker expression on $CD10^+$ neutrophils after coculture with (i)MSC, including representative plots of expression of each marker on neutrophils after culture **B)** Frequencies of $CD10^+$ neutrophils expressing C3AR, CD11b(act), and CD11c, or mean fluorescent intensity (MFI) of CD66b and CD45 in $CD10^+$ neutrophils cultured alone ('neutro only'), in the presence of rhIL-1β, or on iMSC. Lines depict paired samples (n = 4, collected over 3 experiments) **C)** Frequencies of $CD10^+$ neutrophils expressing C3AR, CD11b(act), and CD11c, or mean fluorescent intensity (MFI) of CD66b and CD45 in $CD10^+$ neutrophils cultured alone ('neutro only'), in the presence of MM1S myeloma cells in various quantities, or on iMSC. Lines depict paired samples (n = 4 donors, collected over 3 experiments) **D)** Volcano plot depicting differentially expressed genes of neutrophils cultured on iMSC versus neutrophils cultured alone. $Log_2$FoldChange cutoff, 1; adjusted p-value (padj) cutoff $10^{-4}$. Genes in red are related to inflammation **E)** Enrichment plots for selected Hallmark genesets comparing neutrophils cultured on iMSC to neutrophils cultured alone **F)** BAFF protein in culture supernatant of neutrophils cultured alone ('neutro only'), in the presence of rhIL-1β, on iMSC, or in culture supernatant of (i)MSC cultured without neutrophils ('(i)MSC only') (n = 5 donors, collected over 5 experiments) **G)** BAFF protein in culture supernatant of neutrophils cultured alone ('neutro only') or in the presence of various quantities of MM1S (n = 4 donors, collected over 3 experiments) **H)** IL-1β protein in culture supernatant of neutrophils cultured alone ('neutro only'), on non-inflammatory MSC ('+ MSC'), on iMSC, in the presence of LPS (positive control), or in culture supernatant of iMSC cultured without neutrophils ('iMSC only') Lines depict paired samples (n = 5 donors, collected over 5 experiments) **I)** IL-1β protein in culture supernatant of neutrophils cultured alone ('neutro only') or in the presence of various quantities of MM1S ('+MM1S') (n =4 donors, collected over 3 experiments) **J)** Transcription of *SOCS3* in neutrophils cultured alone ('neutro only'), or in the presence of G-CSF (positive control) and DMSO/Stattic. Lines depict paired samples (n = 5 donors, collected over 3 experiments) **K)** Live cells defined as DAPI-negative in neutrophils cultured alone ('neutro only'), on non-inflammatory MSC ('+MSC'), on iMSC in the presence of DMSO, or on iMSC in the presence of Stattic (n = 5 donors, collected over 5 experiments) **L)** Mean fluorescent intensity (MFI) of CD10 on $CD10^+$ neutrophils on non-inflammatory MSC ('+ MSC'), on iMSC in the presence of DMSO, or on iMSC in the presence of Stattic. Lines depict paired samples (n = 6 donors, collected over 5 experiments).

LPS, lipopolysaccharide. DMSO, dimethyl sulfoxide. Data are presented as mean ± SEM. Significance was calculated in **B**, **C**, **F**, **G**, **H**, **I**, **J**, **K** and **L** using the Wilcoxon Rank Sum test (two-tailed), and in **D** using the Wald test (two-tailed) followed by a Benjamini–Hochberg correction. *P ≤ 0.05, **P ≤ 0.01, NS P > 0.05.

### Supplemental Figure 4

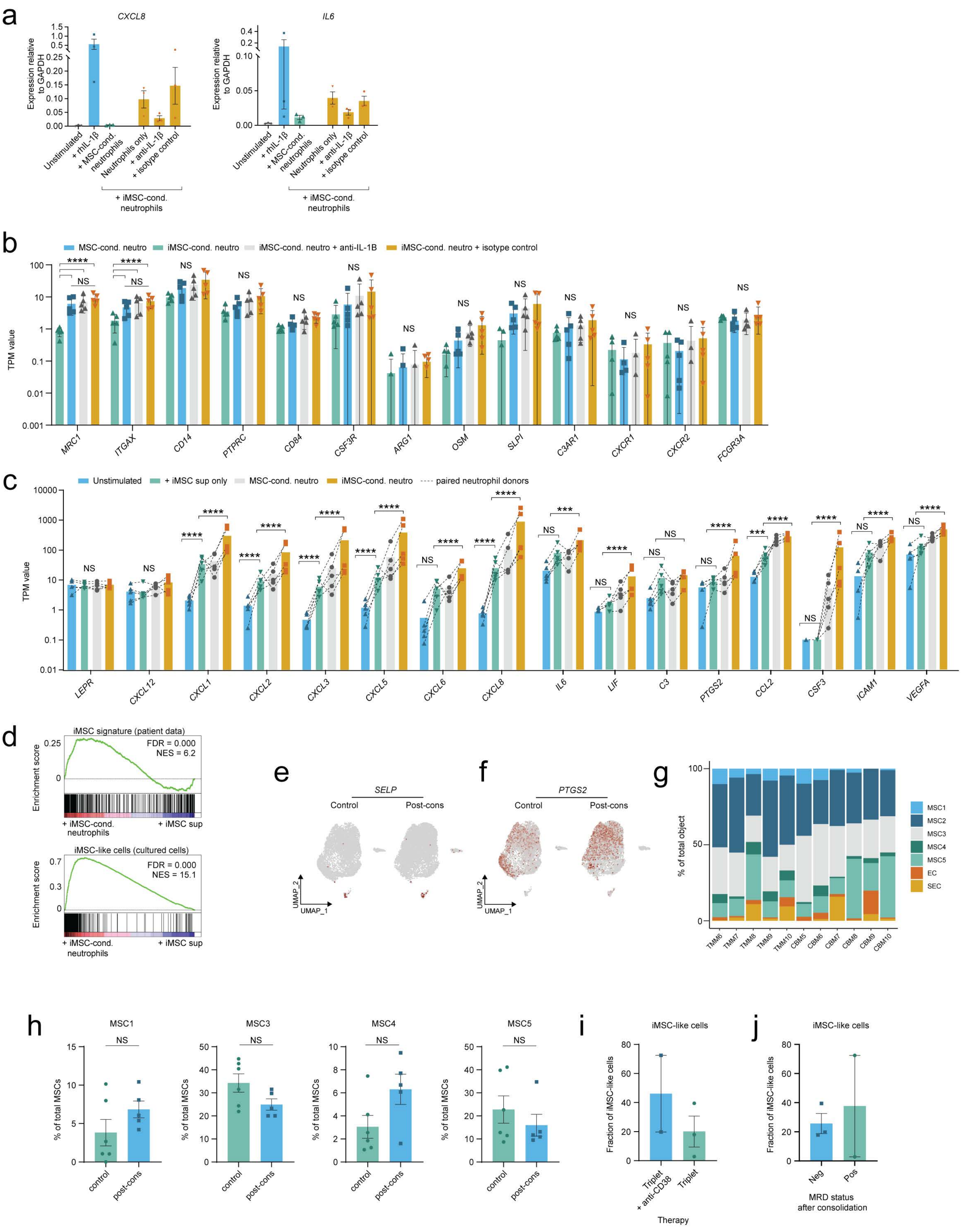

**Supplemental Figure 4** *Related to Figures 4 and 5* **A)** Transcription of *CXCL8* and *IL6* in MSC cultured alone ('unstimulated'), in the presence of rh-IL-1β, in the presence of MSC-conditioned neutrophils, or in the presence of iMSC-conditioned neutrophils with or without anti-IL-1β or an isotype control (n = 3 donors, collected over 3 experiments) **B)** Residual neutrophil transcripts in cultures of MSC that had been cultured with MSC-conditioned neutrophils (blue), iMSC-conditioned neutrophils (green), iMSC-conditioned neutrophils in the presence of anti-IL-1β (grey) or in the presence of an isotype control (orange) (n = 5, collected over 3 experiments) **C)** Transcription of iMSC-related genes in unstimulated MSC (blue), and MSC cultured in the supernatant of iMSC (green), with MSC-conditioned neutrophils (grey) or cultured with iMSC-conditioned neutrophils (orange). Dotted lined depict paired samples (n = 5, collected over 3 experiments) **D)** Enrichment plots for the iMSC-signature (generated from DEGs of single cell RNA sequencing results in [1]), and iMSC-like cells signature (generated from data in Figure 5A) comparing MSC cultured with iMSC-conditioned neutrophils with MSC cultured in supernatant of iMSC. **E)** Transcription of *SELP* in the non-hematopoietic compartment of controls and treated patients **F)** Transcription of *PTGS2* in the non-hematopoietic compartment of controls and treated patients **G)** Cluster distribution in the non-hematopoietic dataset per patient. Colors represent clusters. 'TMM', treated MM-patient. 'CBM', control patient **H)** Percentages of MSCs of total within the discrete MSC clusters per individual in each condition **I)** Expression of 'iMSC-like cell' score per cell in MM patients treated with triplet + anti-CD38 monoclonal therapy or triplet therapy only **J)** Expression of 'iMSC-like cell' score per cell in patients that had become MRD-negative or remained MRD-positive.

MRD, minimal residual disease. Data are presented as mean ± SEM. Significance was calculated in **B** and **C** using the Wald test (two-tailed) followed by a Benjamini–Hochberg correction, and in **H** using the Wilcoxon Rank Sum test (two-tailed). ***P ≤ 0.001, ****P ≤ 0.0001, NS P > 0.05.

### Supplemental Figure 5

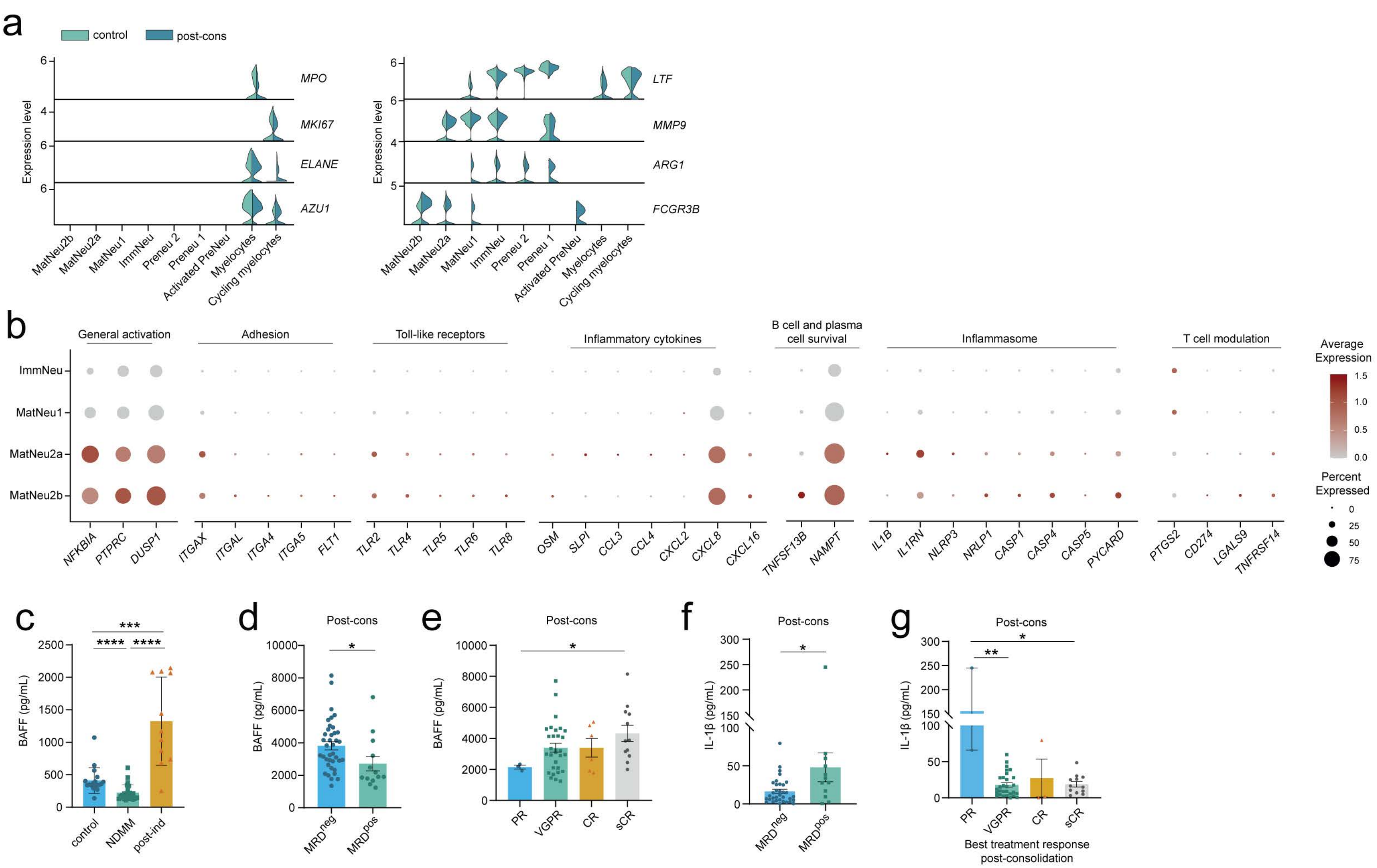

**Supplemental Figure 5** *Related to Figure 6* **A)** Marker gene transcription in the 9 clusters of neutrophilic lineage, split by condition (light green, controls; dark green, treated MM patients) **B)** Transcription of selected genes comparing clusters MatNeu2a and MatNeu2b to clusters MatNeu1 and ImmNeu in treated MM patients **C)** BAFF protein in BM plasma of N = 18 controls, N = 24 NDMM patients and N = 18 patients after induction treatment **D)** BAFF protein in BM plasma of N = 52 patients after consolidation therapy, split by MRD status post-consolidation **E)** BAFF protein in BM plasma of N = 52 patients after consolidation therapy, split by IMWG Uniform Response criteria therapy response[2] **F)** IL-1β protein in BM plasma of N = 52 patients after consolidation therapy, split by MRD status post-consolidation **G)** IL-1β protein in BM plasma of N = 52 patients after consolidation therapy, split by IMWG Uniform Response criteria therapy response[2]

PR, partial response. VGPR, very good partial response. CR, complete response. sCR, stringent complete response. Data are presented as mean ± SEM. Significance was calculated in **C**, **D**, **E**, **F**, and **G** using the Mann–Whitney U test (two-tailed), $^{*}P \leq 0.05$, $^{***}P \leq 0.001$, $^{****}P \leq 0.0001$.

### Supplemental Figure 6

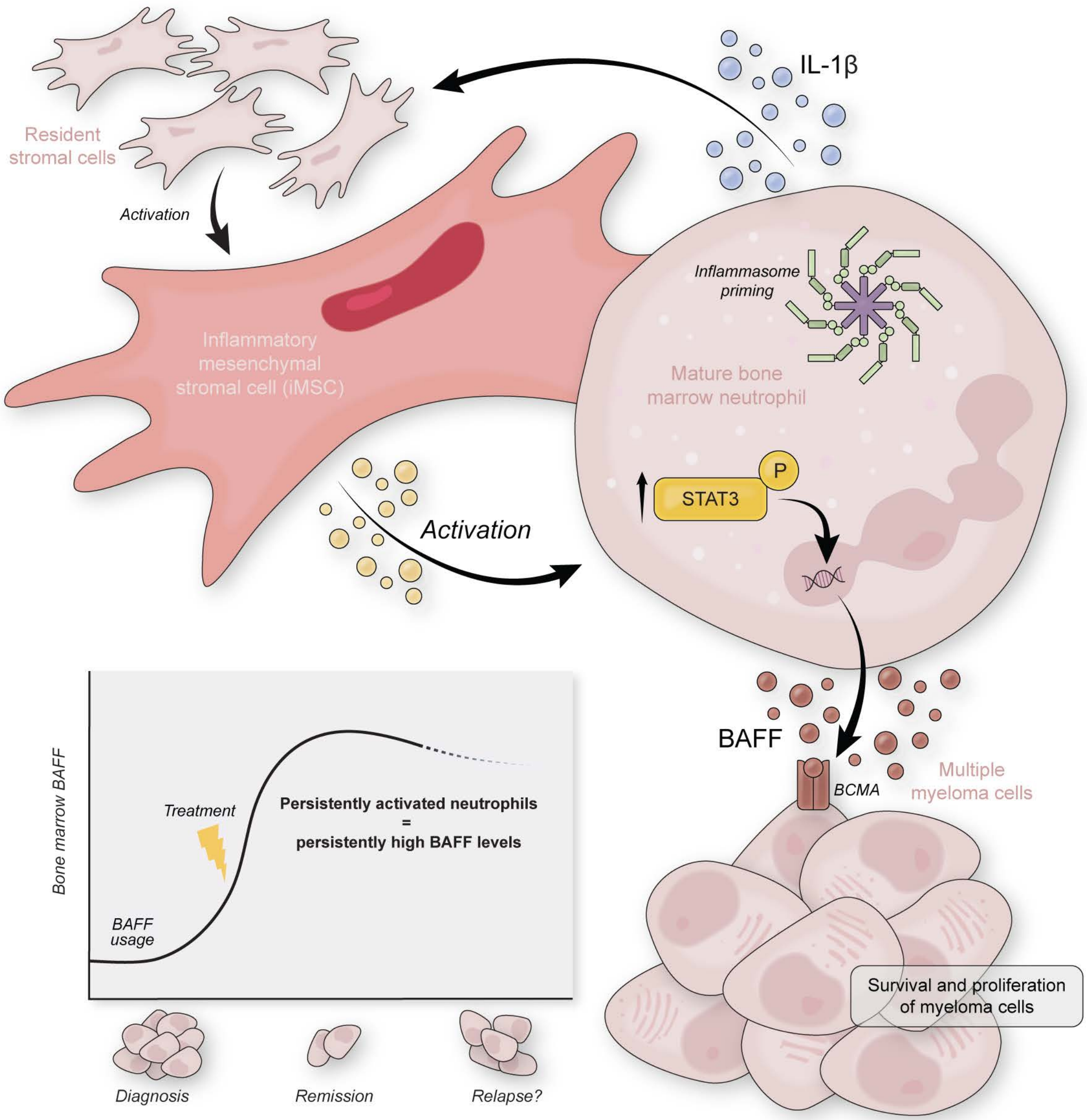

**Supplemental Figure 6** *Model* Interaction with myeloma-specific inflammatory mesenchymal stromal cells (iMSC) leads to an activated phenotype and transcriptome in mature bone marrow neutrophils. Once activated, neutrophils display features of inflammasome priming and secrete IL-1β, which in turn can induce *de novo* stromal inflammation. Moreover, via iMSC-induced STAT3-signaling, activated mature neutrophils produce BAFF, which is a ligand for myeloma-expressed BCMA and induces myeloma cell survival and proliferation. Both iMSC and activated, BAFF-producing neutrophils remain present in the bone marrow of patients after treatment. At diagnosis, when tumor burden is high, BAFF protein is consumed, and detected at low levels in bone marrow plasma. However, when patients are in remission, and tumor cells are mostly gone, continued BAFF production by neutrophils leads to high levels of BAFF protein in the bone marrow, potentially providing residual, therapy-resistant clones with a survival benefit. Therefore, by generating pro-tumor bone marrow inflammation that persists after treatment, neutrophil – stromal cell interactions may impact progression and recurrence of disease.

### Supplemental Table 1

**Supplemental Table 1** Baseline clinical characteristics of NDMM and treated patients included in single cell RNA sequencing analyses of the myeloid and non-hematopoietic BM compartment

| Sample ID | Timepoint of sampling | Treatment | Sex | Age (y)[a] | Genetics[b] | Del17p | Serum M-protein (g/dL) | Plasma cells in BM (%)[c] | Serum rFLC | R-ISS stage | Type of Myeloma | Risk Stratification[d] | MRD[e] after consolidation therapy |
|---|---|---|---|---|---|---|---|---|---|---|---|---|---|
| **MM1** | Diagnosis | N/A | M | 58 | HD | Yes | 3 | 28 | 32.2 | 2 | IgG | High | NEG |
| **MM2** | Diagnosis | N/A | M | 56 | HD | No | 5.6 | 7 | 3.7 | 2 | IgA | Standard | NEG |
| **MM3** | Diagnosis | N/A | F | 68 | t(4;14) | No | 0 | 53 | 201.8 | 3 | FLC | High | NEG |
| **MM4** | Diagnosis | N/A | M | 54 | HD | No | 4.97 | 2 | 47 | 2 | IgG | Standard | NEG |
| **MM5** | Diagnosis | N/A | M | 46 | HD | No | 2.5 | 35 | 10.9 | 1 | IgG | Standard | NEG |
| **MM6** | Diagnosis | N/A | M | 68 | HD | No | 2.69 | N/A | 23.7 | 2 | IgG | Standard | NEG |
| **TMM1** | Post consolidation | triplet | M | 69 | HD | No | 1.37 | 7 | 17.4 | 1 | IgG | Standard | NEG |
| **TMM2** | Post consolidation | triplet | M | 52 | t(11;14) | No | 0 | 1 | 15.1 | 2 | FLC | High | NEG |
| **TMM3** | Post consolidation | triplet | M | 44 | HD | No | 3.07 | 8 | 2.4 | 2 | IgG | Standard | NEG |
| **TMM4** | Post consolidation | triplet + anti-CD38 monoclonal | M | 56 | HD | No | 2.3 | N/A | 174.5 | 1 | IgG | Standard | NEG |
| **TMM5** | Post consolidation | triplet + anti-CD38 monoclonal | M | 53 | t(14;16) | No | 1.34 | 15 | 0.9 | 2 | IgG | High | NEG |
| **TMM6** | Post consolidation | triplet | F | 65 | HD | No | 3.32 | 30 | 50.9 | 2 | IgG | Standard | POS |
| **TMM7** | Post consolidation | triplet | M | 64 | HD | No | 2.84 | 12 | 0.9 | 1 | IgG | Standard | NEG |
| **TMM8** | Post consolidation | triplet + anti-CD38 monoclonal | M | 47 | HD | No | 3.9 | 40 | 4.79 | 3 | IgG | Standard | POS |
| **TMM9** | Post consolidation | triplet + anti-CD38 monoclonal | F | 52 | HD | No | 4.62 | 52 | 0.008 | 2 | IgG | Standard | NEG |
| **TMM10** | Post consolidation | triplet + anti-CD38 monoclonal | M | 59 | HD | No | 2.13 | 50 | 1.31 | 1 | IgG | Standard | NEG |

**a** at time of tissue collection

**b** HD; hyperdiploidy, t(xx;xx); translocation

**c** as determined by $CD38^+$/$CD138^+$ flow cytometry in bone marrow aspirates

**d** based on FISH, the presence of del17p or t(4;14)

**e** determined by flow cytometry, cut-off at $10^{-5}$

KRd; carfilzomib, lenalidomide, dexamethasone, VTD; bortezomib, thalidomide, dexamethasone, D-VTD; daratumumab, bortezomib, thalidomide, dexamethasone

N/A; not applicable/available, FLC; free light chain, MRD; minimal residual disease, (y) years, (d) days

### Supplemental Table 2

**Supplemental Table 2** Clinical characteristics of non-cancer controls included in single cell RNA sequencing analyses of the myeloid and the non-hematopoietic BM compartment

| Sample ID | Sex | Age (y)[a] | Source |
|---|---|---|---|
| **CBM1** | M | 65 | Sternum |
| **CBM2** | F | 71 | Sternum |
| **CBM3** | M | 61 | Sternum |
| **CBM4** | F | 59 | Sternum |
| **CBM5** | M | 70 | Sternum |
| **CBM6** | M | 27 | Sternum |
| **CBM7** | M | 53 | Sternum |
| **CBM8** | M | 75 | Femur head |
| **CBM9** | F | 57 | Femur head |
| **CBM10** | M | 72 | Femur head |

[a] at time of tissue collection

### Supplemental Table 3

**Supplemental Table 3** Results of gene set enrichment analysis (GSEA) of HALLMARK pathways comparing cluster MatNeu2 of NDMM with that of controls

| Geneset | Size | NES | Nominal p-value | FDR q-value[a] |
|---|---|---|---|---|
| **Up in NDMM versus controls** | | | | |
| HALLMARK_INTERFERON_GAMMA_RESPONSE | 197 | 7.15 | 0.000 | 0.000 |
| HALLMARK_INTERFERON_ALPHA_RESPONSE | 95 | 6.97 | 0.000 | 0.000 |
| HALLMARK_TNFA_SIGNALING_VIA_NFKB | 198 | 5.08 | 0.000 | 0.000 |
| HALLMARK_INFLAMMATORY_RESPONSE | 183 | 4.51 | 0.000 | 0.000 |
| HALLMARK_IL6_JAK_STAT3_SIGNALING | 84 | 3.57 | 0.000 | 0.000 |
| HALLMARK_ALLOGRAFT_REJECTION | 188 | 3.20 | 0.000 | 0.000 |
| HALLMARK_HYPOXIA | 182 | 2.68 | 0.000 | 0.000 |
| HALLMARK_COMPLEMENT | 189 | 2.66 | 0.000 | 0.000 |
| HALLMARK_PI3K_AKT_MTOR_SIGNALING | 98 | 2.54 | 0.000 | 0.000 |
| HALLMARK_IL2_STAT5_SIGNALING | 193 | 2.49 | 0.000 | 0.000 |
| HALLMARK_KRAS_SIGNALING_DN | 129 | 2.37 | 0.000 | 0.001 |
| HALLMARK_COAGULATION | 115 | 2.29 | 0.000 | 0.002 |
| HALLMARK_APOPTOSIS | 158 | 2.20 | 0.000 | 0.003 |
| HALLMARK_EPITHELIAL_MESENCHYMAL_TRANSITION | 178 | 2.15 | 0.000 | 0.004 |
| HALLMARK_UV_RESPONSE_UP | 151 | 2.10 | 0.000 | 0.005 |
| HALLMARK_KRAS_SIGNALING_UP | 175 | 1.82 | 0.012 | 0.025 |
| HALLMARK_XENOBIOTIC_METABOLISM | 172 | 1.72 | 0.027 | 0.041 |
| **Down in NDMM versus controls** | | | | |
| HALLMARK_MYC_TARGETS_V1 | 194 | -8.01 | 0.000 | 0.000 |
| HALLMARK_OXIDATIVE_PHOSPHORYLATION | 183 | -6.42 | 0.000 | 0.000 |
| HALLMARK_DNA_REPAIR | 147 | -4.07 | 0.000 | 0.000 |
| HALLMARK_UNFOLDED_PROTEIN_RESPONSE | 107 | -4.07 | 0.000 | 0.000 |
| HALLMARK_E2F_TARGETS | 195 | -3.84 | 0.000 | 0.000 |
| HALLMARK_PROTEIN_SECRETION | 93 | -3.74 | 0.000 | 0.000 |
| HALLMARK_MTORC1_SIGNALING | 196 | -3.70 | 0.000 | 0.000 |
| HALLMARK_ADIPOGENESIS | 185 | -3.40 | 0.000 | 0.000 |
| HALLMARK_MYC_TARGETS_V2 | 58 | -3.19 | 0.000 | 0.000 |
| HALLMARK_P53_PATHWAY | 187 | -3.18 | 0.000 | 0.000 |
| HALLMARK_G2M_CHECKPOINT | 189 | -2.96 | 0.000 | 0.000 |
| HALLMARK_FATTY_ACID_METABOLISM | 145 | -2.90 | 0.000 | 0.000 |
| HALLMARK_MITOTIC_SPINDLE | 197 | -2.71 | 0.002 | 0.000 |
| HALLMARK_CHOLESTEROL_HOMEOSTASIS | 70 | -2.57 | 0.000 | 0.000 |
| HALLMARK_PEROXISOME | 95 | -2.12 | 0.004 | 0.004 |
| HALLMARK_HEME_METABOLISM | 189 | -2.01 | 0.004 | 0.007 |
| HALLMARK_REACTIVE_OXYGEN_SPECIES_PATHWAY | 47 | -1.81 | 0.023 | 0.020 |
| HALLMARK_ANDROGEN_RESPONSE | 93 | -1.72 | 0.028 | 0.029 |

[a] Gene sets were considered significantly up-/downregulated when FDR < 0.05
NES; Normalized Enrichment Score, FDR; false discovery rate

### Supplemental Table 4

**Supplemental Table 4** Top 50 differentially expressed genes (DEGs) comparing cluster MatNeu2 of NDMM to that of controls

| Gene name | Average Log2FC | Adj. p-value[a] | Gene name | Average Log2FC | Adj. p-value[a] |
|---|---|---|---|---|---|
| **Up in NDMM versus controls** | | | **Down in NDMM versus controls** | | |
| MNDA | 0.92 | 0 | RPS29 | -0.82 | 3.53E-245 |
| IER3 | 0.87 | 3.75E-204 | RPS24 | -0.81 | 1.36E-241 |
| IFIT3 | 0.84 | 7.19E-143 | RPL9 | -0.74 | 1.72E-249 |
| NFKBIA | 0.83 | 2.65E-275 | RPL21 | -0.73 | 6.69E-209 |
| PI3 | 0.82 | 1.09E-126 | RPL34 | -0.71 | 1.22E-289 |
| IL1RN | 0.82 | 4.79E-167 | RPS28 | -0.69 | 3.91E-177 |
| TNFAIP3 | 0.81 | 1.99E-180 | VCAN | -0.68 | 8.01E-245 |
| IFITM3 | 0.80 | 5.05E-161 | RPS27 | -0.68 | 6.00E-250 |
| IRF1 | 0.79 | 2.23E-178 | RPL11 | -0.68 | 1.91E-187 |
| FCGR3B | 0.77 | 3.27E-199 | RPL32 | -0.66 | 1.73E-167 |
| FFAR2 | 0.71 | 2.30E-135 | RPS6 | -0.66 | 4.65E-170 |
| PPP1R15A | 0.71 | 6.64E-159 | RPS8 | -0.65 | 3.69E-148 |
| CST7 | 0.70 | 3.58E-136 | RPL38 | -0.64 | 1.47E-162 |
| MSRB1 | 0.70 | 1.70E-140 | RPL35A | -0.63 | 7.70E-160 |
| MARCKS | 0.68 | 4.39E-167 | RPL6 | -0.62 | 1.43E-151 |
| IFIT2 | 0.68 | 1.20E-104 | RPLP2 | -0.61 | 6.19E-147 |
| FAM65B | 0.66 | 1.41E-124 | RPL37 | -0.61 | 8.71E-217 |
| PARP14 | 0.65 | 5.64E-112 | MT-CO1 | -0.61 | 9.07E-156 |
| MYADM | 0.65 | 1.79E-127 | RPL26 | -0.61 | 3.99E-149 |
| FCER1G | 0.62 | 4.92E-156 | MT-ND3 | -0.60 | 2.72E-177 |
| TNFAIP6 | 0.60 | 5.62E-95 | RPL39 | -0.60 | 0 |
| IFITM2 | 0.60 | 0 | RPL37A | -0.59 | 1.55E-153 |
| IFITM1 | 0.59 | 3.31E-102 | EEF1A1 | -0.59 | 6.87E-135 |
| TSC22D3 | 0.59 | 3.97E-112 | MT-CO3 | -0.58 | 2.72E-224 |
| SMCHD1 | 0.59 | 3.86E-196 | RIOK3 | -0.57 | 8.38E-138 |
| AQP9 | 0.58 | 1.90E-117 | PTMA | -0.57 | 8.56E-121 |
| ISG15 | 0.58 | 3.78E-81 | RPL12 | -0.57 | 2.80E-120 |
| TAGLN2 | 0.57 | 3.51E-119 | FAM101B | -0.56 | 1.85E-139 |
| FOSB | 0.57 | 5.31E-93 | HMGB1 | -0.56 | 3.95E-141 |
| ADGRE5 | 0.57 | 4.53E-95 | RPS13 | -0.56 | 4.25E-118 |
| ALPL | 0.56 | 2.26E-98 | SYF2 | -0.55 | 2.07E-125 |
| CXCR2 | 0.55 | 6.93E-91 | RPL36 | -0.55 | 9.95E-108 |
| PRELID1 | 0.55 | 4.12E-85 | FCN1 | -0.55 | 2.11E-125 |
| CLEC4E | 0.54 | 7.27E-86 | RPL22 | -0.53 | 1.70E-145 |
| IFIT1 | 0.54 | 3.47E-83 | RPL24 | -0.52 | 8.01E-106 |
| RILPL2 | 0.54 | 6.98E-86 | RPS7 | -0.51 | 1.33E-108 |
| PFN1 | 0.54 | 6.14E-153 | RPS15A | -0.51 | 1.15E-136 |
| SLC38A2 | 0.53 | 1.26E-83 | RPS21 | -0.51 | 6.67E-108 |
| FAM129A | 0.53 | 3.94E-82 | IRS2 | -0.50 | 1.42E-103 |
| IFI16 | 0.52 | 3.01E-84 | RPS25 | -0.50 | 2.61E-101 |
| CLIC1 | 0.52 | 1.28E-95 | RPS3A | -0.48 | 2.42E-95 |
| FYB | 0.52 | 2.83E-83 | RPL23 | -0.48 | 6.31E-100 |
| NFKBIZ | 0.51 | 3.46E-84 | RPL5 | -0.48 | 7.82E-112 |
| MAPK14 | 0.51 | 4.12E-77 | NACA | -0.48 | 8.49E-96 |
| SOD2 | 0.50 | 0 | IRF2BP2 | -0.48 | 9.07E-90 |
| MX2 | 0.50 | 4.12E-76 | RPL35 | -0.47 | 4.77E-97 |
| MX1 | 0.49 | 2.57E-80 | HIF1A | -0.47 | 1.40E-150 |
| ISG20 | 0.48 | 9.35E-74 | MAFB | -0.47 | 2.07E-173 |
| TAP1 | 0.48 | 2.43E-77 | RPL19 | -0.46 | 1.02E-80 |
| LDHA | 0.47 | 5.29E-59 | RPL14 | -0.46 | 1.73E-91 |

Computed using Seurat's FindMarkers() function

[a] Differential gene expression considered significant when adj. p-value cutoff $< 10^{-4}$

Log2FC; Log2 fold change, Adj. p-value; adjusted p-value

### Supplemental Table 5

**Supplemental Table 5** Results of gene set enrichment analysis (GSEA) of HALLMARK pathways comparing clusters MatNeu2 and MatNeu1 in NDMM

| Geneset | Size | NES | Nominal p-value | FDR q-value[a] |
|---|---|---|---|---|
| **Up in MatNeu2 versus MatNeu1** | | | | |
| HALLMARK_TNFA_SIGNALING_VIA_NFKB | 198 | 8.16 | 0.000 | 0.000 |
| HALLMARK_INTERFERON_GAMMA_RESPONSE | 197 | 7.96 | 0.000 | 0.000 |
| HALLMARK_INTERFERON_ALPHA_RESPONSE | 95 | 7.30 | 0.000 | 0.000 |
| HALLMARK_INFLAMMATORY_RESPONSE | 183 | 5.39 | 0.000 | 0.000 |
| HALLMARK_IL6_JAK_STAT3_SIGNALING | 84 | 4.34 | 0.000 | 0.000 |
| HALLMARK_APOPTOSIS | 158 | 4.11 | 0.000 | 0.000 |
| HALLMARK_PI3K_AKT_MTOR_SIGNALING | 98 | 3.36 | 0.000 | 0.000 |
| HALLMARK_UV_RESPONSE_UP | 151 | 3.27 | 0.000 | 0.000 |
| HALLMARK_P53_PATHWAY | 187 | 3.19 | 0.000 | 0.000 |
| HALLMARK_ALLOGRAFT_REJECTION | 188 | 3.14 | 0.000 | 0.000 |
| HALLMARK_IL2_STAT5_SIGNALING | 193 | 3.01 | 0.000 | 0.000 |
| HALLMARK_HYPOXIA | 182 | 3.01 | 0.000 | 0.000 |
| HALLMARK_COMPLEMENT | 189 | 2.94 | 0.000 | 0.000 |
| HALLMARK_MTORC1_SIGNALING | 196 | 2.90 | 0.000 | 0.000 |
| HALLMARK_UNFOLDED_PROTEIN_RESPONSE | 107 | 2.78 | 0.000 | 0.000 |
| HALLMARK_ANDROGEN_RESPONSE | 93 | 2.51 | 0.000 | 0.000 |
| HALLMARK_KRAS_SIGNALING_UP | 175 | 2.15 | 0.000 | 0.004 |
| HALLMARK_MYOGENESIS | 162 | 1.95 | 0.006 | 0.012 |
| HALLMARK_XENOBIOTIC_METABOLISM | 172 | 1.92 | 0.006 | 0.013 |
| HALLMARK_ESTROGEN_RESPONSE_EARLY | 179 | 1.85 | 0.013 | 0.018 |
| HALLMARK_TGF_BETA_SIGNALING | 54 | 1.85 | 0.014 | 0.017 |
| HALLMARK_ESTROGEN_RESPONSE_LATE | 175 | 1.84 | 0.012 | 0.018 |
| **Down in MatNeu2 versus MatNeu1** | | | | |
| HALLMARK_OXIDATIVE_PHOSPHORYLATION | 183 | -4.82 | 0.000 | 0.000 |
| HALLMARK_MYC_TARGETS_V1 | 194 | -3.99 | 0.000 | 0.000 |
| HALLMARK_MITOTIC_SPINDLE | 197 | -3.21 | 0.000 | 0.000 |
| HALLMARK_G2M_CHECKPOINT | 189 | -2.91 | 0.000 | 0.000 |
| HALLMARK_ADIPOGENESIS | 185 | -2.65 | 0.000 | 0.000 |
| HALLMARK_REACTIVE_OXYGEN_SPECIES_PATHWAY | 47 | -2.34 | 0.002 | 0.002 |
| HALLMARK_E2F_TARGETS | 195 | -2.33 | 0.000 | 0.002 |
| HALLMARK_GLYCOLYSIS | 187 | -2.23 | 0.000 | 0.003 |
| HALLMARK_FATTY_ACID_METABOLISM | 145 | -2.21 | 0.000 | 0.003 |
| HALLMARK_PROTEIN_SECRETION | 93 | -2.20 | 0.008 | 0.003 |
| HALLMARK_DNA_REPAIR | 147 | -2.18 | 0.002 | 0.003 |
| HALLMARK_PEROXISOME | 95 | -2.10 | 0.000 | 0.005 |
| HALLMARK_MYC_TARGETS_V2 | 58 | -1.74 | 0.025 | 0.033 |

[a] Gene sets were considered significantly up-/downregulated when FDR < 0.05
NES; Normalized Enrichment Score, FDR; false discovery rate

### Supplemental Table 6

**Supplemental Table 6** Top 50 differentially expressed genes (DEGs) comparing clusters MatNeu2 and MatNeu1 in NDMM

| Gene name | Average Log2FC | Adj. p-value[a] | Gene name | Average Log2FC | Adj. p-value[a] |
|---|---|---|---|---|---|
| Up in MatNeu2 versus MatNeu1 | | | Down in MatNeu2 versus MatNeu1 | | |
| IFITM3 | 0.75 | 3.01E-146 | MMP9 | -1.10 | 0 |
| AIF1 | 0.68 | 2.39E-119 | S100A12 | -0.81 | 0 |
| IFIT2 | 0.62 | 7.01E-82 | FAM101B | -0.79 | 9.53E-217 |
| SLC25A37 | 0.58 | 1.40E-162 | HMGB2 | -0.72 | 7.57E-175 |
| CLEC7A | 0.57 | 3.23E-95 | PGLYRP1 | -0.68 | 3.88E-159 |
| ZFP36L1 | 0.54 | 1.34E-95 | PADI4 | -0.66 | 4.86E-158 |
| IFIT3 | 0.52 | 2.37E-56 | CDA | -0.60 | 2.08E-154 |
| TNFAIP2 | 0.51 | 4.44E-72 | EMB | -0.50 | 7.14E-88 |
| GBP2 | 0.50 | 2.88E-67 | ARG1 | -0.50 | 6.49E-84 |
| SEC14L1 | 0.47 | 3.15E-70 | FCN1 | -0.50 | 3.67E-100 |
| GADD45B | 0.46 | 3.30E-47 | TSPO | -0.46 | 5.67E-129 |
| PLK3 | 0.46 | 1.25E-62 | ALOX5AP | -0.46 | 1.34E-102 |
| FAM129A | 0.46 | 3.13E-60 | CYBB | -0.45 | 4.86E-78 |
| CXCR2 | 0.46 | 3.15E-56 | VIM | -0.44 | 2.20E-112 |
| ADGRE5 | 0.46 | 9.38E-61 | C9orf16 | -0.42 | 8.73E-69 |
| NEAT1 | 0.46 | 0 | TXN | -0.42 | 3.19E-73 |
| PI3 | 0.44 | 2.41E-30 | S100P | -0.41 | 9.57E-133 |
| TAGLN2 | 0.43 | 1.02E-75 | PGD | -0.41 | 8.62E-59 |
| IFITM1 | 0.43 | 2.71E-52 | LGALS1 | -0.41 | 5.58E-54 |
| AQP9 | 0.43 | 6.22E-72 | CALM1 | -0.40 | 2.20E-59 |
| FFAR2 | 0.42 | 4.36E-48 | RIOK3 | -0.40 | 5.03E-52 |
| MX2 | 0.42 | 3.02E-49 | PYGL | -0.39 | 1.67E-52 |
| CEBPB | 0.42 | 6.90E-81 | RPS15A | -0.39 | 1.27E-54 |
| NINJ1 | 0.41 | 6.77E-70 | PPP2R5A | -0.38 | 6.16E-51 |
| VMP1 | 0.40 | 3.13E-50 | TKT | -0.38 | 1.96E-56 |
| ISG15 | 0.40 | 5.68E-33 | CTSD | -0.38 | 2.13E-49 |
| TREM1 | 0.40 | 1.79E-47 | RPL27 | -0.37 | 5.81E-50 |
| IFIT1 | 0.39 | 6.06E-35 | SLC37A3 | -0.37 | 2.63E-66 |
| RNF213 | 0.39 | 1.11E-39 | ITGAM | -0.37 | 1.54E-50 |
| FCGR2A | 0.39 | 9.07E-43 | ANXA1 | -0.37 | 1.84E-43 |
| KDM6B | 0.39 | 5.36E-44 | SYNE1 | -0.37 | 1.17E-53 |
| SLC15A4 | 0.38 | 3.48E-41 | CTB-61M7.2 | -0.36 | 1.63E-44 |
| ADGRE2 | 0.38 | 6.73E-55 | RGL4 | -0.35 | 1.06E-45 |
| PARP14 | 0.38 | 8.22E-35 | GYG1 | -0.35 | 2.15E-50 |
| CLEC2B | 0.37 | 1.11E-35 | RPL39 | -0.35 | 6.67E-81 |
| TLR2 | 0.37 | 8.93E-40 | RBP7 | -0.34 | 4.30E-49 |
| ITGAX | 0.37 | 5.43E-39 | S100A4 | -0.33 | 4.87E-166 |
| LINC01272 | 0.37 | 8.73E-38 | CRISP3 | -0.33 | 9.36E-66 |
| MX1 | 0.36 | 1.99E-34 | CSTA | -0.32 | 1.72E-37 |
| ZFP36 | 0.35 | 4.54E-49 | GAPDH | -0.32 | 2.08E-63 |
| G0S2 | 0.35 | 1.64E-75 | METTL9 | -0.31 | 1.50E-30 |
| AMPD2 | 0.35 | 4.20E-33 | S100A8 | -0.31 | 0 |
| SQSTM1 | 0.35 | 7.68E-32 | MYL6 | -0.31 | 6.75E-96 |
| GBP5 | 0.34 | 6.12E-37 | CAP1 | -0.31 | 1.93E-47 |
| HERC5 | 0.34 | 3.65E-35 | CAPG | -0.31 | 7.16E-46 |
| PPP1R15A | 0.34 | 6.15E-44 | CKAP4 | -0.30 | 1.06E-30 |
| MARCKS | 0.34 | 2.98E-58 | MEGF9 | -0.30 | 3.04E-29 |
| RP11-22N19.2 | 0.34 | 1.88E-44 | TUBA4A | -0.30 | 1.01E-29 |
| ICAM1 | 0.33 | 1.44E-30 | C14orf2 | -0.29 | 2.83E-28 |
| RNASET2 | 0.33 | 9.30E-32 | MYL12A | -0.29 | 3.23E-35 |

Computed using Seurat's FindMarkers() function

[a] Differential gene expression considered significant when adj. p-value cutoff < $10^{-4}$

Log2FC; Log2 fold change, Adj. p-value; adjusted p-value

### Supplemental Table 7

**Supplemental Table 7** Top 50 genes up- and downregulated in ADSC stimulated with rhIL-1β compared to unstimulated ADSC

| Gene name | Average Log2FC | Adj. p-value[a] | Gene name | Average Log2FC | Adj. p-value[a] |
|---|---|---|---|---|---|
| **Up when stimulated with rhIL-1β** | | | **Down when stimulated with rhIL-1β** | | |
| CSF3 | 16.91 | 2.14E-46 | ENSG00000288534 | -5.67 | 4.29E-07 |
| CXCL8 | 13.91 | 1.64E-152 | ADH1B | -5.64 | 3.80E-07 |
| CSF2 | 12.45 | 3.25E-25 | ACTC1 | -5.34 | 3.27E-36 |
| CXCL3 | 11.08 | 3.42E-125 | HTR2B | -4.89 | 2.62E-27 |
| CXCL1 | 10.57 | 2.65E-237 | RGS4 | -4.47 | 3.41E-28 |
| CXCL6 | 10.55 | 3.22E-52 | OXTR | -4.44 | 3.14E-12 |
| SERPINB4 | 10.52 | 3.50E-18 | TPD52L1 | -4.01 | 6.30E-25 |
| CXCL5 | 10.24 | 8.41E-145 | ADRA2C | -3.96 | 2.25E-11 |
| IL36RN | 10.14 | 4.86E-33 | SAMD11 | -3.80 | 2.28E-08 |
| CXCL2 | 9.06 | 2.70E-80 | KRT14 | -3.72 | 3.45E-14 |
| CCL20 | 8.51 | 1.84E-25 | FHL1 | -3.71 | 4.24E-12 |
| IL1B | 7.86 | 1.55E-43 | PLCL1 | -3.60 | 1.08E-10 |
| ADORA2A | 7.74 | 1.45E-10 | MYH2 | -3.57 | 4.86E-09 |
| IL1RN | 7.39 | 1.11E-60 | RIPOR2 | -3.42 | 8.73E-06 |
| PPP4R4 | 7.30 | 9.00E-15 | IRAG1 | -3.37 | 7.71E-05 |
| DCLK3 | 6.53 | 1.67E-25 | NPR3 | -3.28 | 5.34E-21 |
| IL24 | 6.35 | 2.18E-17 | DNM3 | -3.01 | 1.07E-07 |
| CCL2 | 6.21 | 4.17E-41 | ASPN | -2.98 | 2.83E-12 |
| SERPINA1 | 6.09 | 2.66E-19 | CCN3 | -2.93 | 2.87E-10 |
| RHCG | 5.99 | 1.62E-11 | SCN9A | -2.91 | 9.21E-20 |
| LIF | 5.91 | 3.54E-23 | KRT18 | -2.89 | 1.22E-13 |
| IL33 | 5.71 | 1.57E-31 | CRYAB | -2.86 | 1.27E-08 |
| IL6 | 5.61 | 1.38E-66 | LMOD1 | -2.85 | 1.29E-10 |
| SOD2 | 5.49 | 1.64E-45 | C11orf87 | -2.84 | 1.20E-10 |
| MMP8 | 5.31 | 1.87E-15 | TRIB2 | -2.80 | 6.24E-11 |
| LRRN3 | 5.16 | 3.06E-29 | SAMD3 | -2.78 | 1.10E-05 |
| PTGS2 | 5.12 | 2.95E-17 | MALL | -2.76 | 8.95E-09 |
| CD82 | 4.94 | 4.04E-38 | SYNDIG1 | -2.74 | 7.15E-10 |
| SERPINB2 | 4.94 | 1.22E-40 | MAN1C1 | -2.69 | 3.27E-12 |
| HSD11B1 | 4.93 | 9.51E-10 | RHOJ | -2.69 | 9.05E-09 |
| NFATC2 | 4.87 | 2.51E-10 | SGCG | -2.66 | 3.46E-09 |
| SLAMF8 | 4.86 | 4.45E-16 | SCUBE3 | -2.65 | 8.53E-24 |
| KCNN3 | 4.83 | 1.44E-17 | LMO3 | -2.63 | 2.75E-05 |
| ICAM1 | 4.82 | 7.82E-10 | VIT | -2.58 | 3.85E-06 |
| MMP10 | 4.78 | 1.00E-09 | ADGRF5 | -2.58 | 4.23E-05 |
| POU2F2 | 4.72 | 1.25E-09 | MYOZ2 | -2.57 | 1.23E-06 |
| NFKBIZ | 4.70 | 5.37E-13 | KCND3 | -2.57 | 1.97E-12 |
| TMEM132B | 4.63 | 1.15E-36 | SLC38A4 | -2.55 | 3.51E-09 |
| GRIA1 | 4.62 | 3.74E-13 | NR2F2 | -2.49 | 1.05E-17 |
| EDNRB | 4.57 | 4.60E-12 | AKR1B10 | -2.48 | 2.27E-06 |
| TFPI2 | 4.47 | 5.88E-29 | LDB2 | -2.47 | 1.76E-13 |
| NTN1 | 4.45 | 2.01E-14 | AK5 | -2.45 | 9.98E-13 |
| CLDN1 | 4.39 | 6.54E-16 | CLEC14A | -2.45 | 9.99E-09 |
| RRAD | 4.35 | 4.00E-24 | LBH | -2.44 | 1.29E-08 |
| MMP1 | 4.34 | 2.38E-21 | KCNS2 | -2.44 | 0.000399 |
| C3 | 4.34 | 2.91E-31 | ENSG00000277758 | -2.41 | 7.09E-14 |
| ZC3H12A | 4.31 | 6.61E-11 | RCAN2 | -2.41 | 1.59E-06 |
| TRIM36 | 4.25 | 6.36E-15 | SLC16A14 | -2.40 | 1.73E-07 |
| AMPD3 | 4.24 | 1.46E-07 | SSTR1 | -2.40 | 3.37E-06 |
| MMP12 | 4.23 | 2.66E-16 | CLIC3 | -2.37 | 4.79E-11 |

Computed using DESeq2. Significance was calculated using the Wald test (two-tailed) followed by the Benjamini-Hochberg correction

[a] Differential gene expression considered significant when adj. p-value cutoff < $10^{-4}$

Log2FC; Log2 fold change, Adj. p-value; adjusted p-value

### Supplemental Table 8

**Supplemental Table 8** Top 50 genes up- and downregulated in neutrophils cultured on stroma compared to cultured alone

| Gene name | Average Log2FC | Adj. p-value[a] | Gene name | Average Log2FC | Adj. p-value[a] |
|---|---|---|---|---|---|
| **Up when cultured on stroma** | | | **Down when cultured on stroma** | | |
| MAP1B | 7.08 | 9.66E-46 | INTS3 | -33.74 | 1.18E-05 |
| COL1A2 | 8.08 | 1.63E-32 | PPP1R11 | -6.60 | 0.000961 |
| MMP3 | 10.10 | 1.29E-28 | | | |
| COL1A1 | 7.11 | 1.82E-26 | | | |
| EHD2 | 5.62 | 6.15E-26 | | | |
| LOX | 8.69 | 3.32E-24 | | | |
| HAS2 | 8.44 | 1.47E-21 | | | |
| GNG12 | 5.42 | 2.52E-21 | | | |
| THY1 | 7.11 | 5.13E-21 | | | |
| FSTL1 | 6.40 | 1.06E-20 | | | |
| COL3A1 | 11.61 | 2.52E-17 | | | |
| MMP2 | 4.27 | 3.17E-17 | | | |
| FLNC | 6.64 | 7.68E-17 | | | |
| CAV1 | 4.49 | 1.66E-16 | | | |
| FN1 | 6.13 | 1.73E-16 | | | |
| COL6A1 | 3.03 | 2.32E-15 | | | |
| CAVIN1 | 4.63 | 2.32E-15 | | | |
| S100A16 | 5.88 | 2.68E-14 | | | |
| CCN1 | 9.67 | 3.23E-14 | | | |
| TFPI2 | 5.35 | 5.03E-13 | | | |
| CLMP | 4.07 | 6.91E-13 | | | |
| COL6A3 | 4.13 | 9.11E-13 | | | |
| ENPP2 | 1.84 | 2.05E-12 | | | |
| CCDC80 | 4.23 | 8.75E-12 | | | |
| DKK1 | 9.97 | 9.89E-12 | | | |
| POSTN | 5.55 | 2.22E-11 | | | |
| TMEM47 | 6.43 | 3.40E-11 | | | |
| SERPINE1 | 4.49 | 1.16E-10 | | | |
| SOCS3 | 1.62 | 1.57E-10 | | | |
| SDC1 | 3.85 | 1.58E-10 | | | |
| PRRX1 | 8.33 | 3.46E-10 | | | |
| PODXL | 3.01 | 3.52E-10 | | | |
| RAB3B | 4.14 | 3.53E-10 | | | |
| KDELR3 | 3.82 | 3.58E-10 | | | |
| OLFML3 | 5.15 | 3.99E-10 | | | |
| FGF2 | 5.23 | 4.53E-10 | | | |
| RND3 | 6.20 | 6.00E-10 | | | |
| MXRA8 | 3.15 | 7.11E-10 | | | |
| CDH2 | 3.89 | 7.11E-10 | | | |
| CDC20 | 2.58 | 9.57E-10 | | | |
| MMP1 | 10.27 | 1.93E-09 | | | |
| SNAI2 | 8.42 | 3.03E-09 | | | |
| CREB3L1 | 4.75 | 6.18E-09 | | | |
| FRMD6 | 2.62 | 9.36E-09 | | | |
| AXL | 2.89 | 1.13E-08 | | | |
| PLS3 | 4.64 | 1.61E-08 | | | |
| TUBB3 | 3.41 | 1.69E-08 | | | |
| LOXL2 | 2.33 | 2.02E-08 | | | |
| SCARA3 | 7.45 | 4.13E-08 | | | |
| SMTN | 1.70 | 5.89E-08 | | | |

Computed using DESeq2. Significance was calculated using the Wald test (two-tailed) followed by the Benjamini-Hochberg correction

[a] Differential gene expression considered significant when adj. p-value cutoff < $10^{-4}$

Log2FC; Log2 fold change, Adj. p-value; adjusted p-value

### Supplemental Table 9

**Supplemental Table 9** Top 50 genes up- and downregulated in neutrophils cultured on iMSC compared to neutrophils cultured on MSC

| Gene name | Average Log2FC | Adj. p-value[a] | Gene name | Average Log2FC | Adj. p-value[a] |
|---|---|---|---|---|---|
| Up when cultured on iMSC | | | Down when cultured on iMSC | | |
| NLRP2 | 26.05 | 3.2162E-05 | FAM120B | -34.86 | 3.1466E-10 |
| INTS3 | 23.57 | 0.00141968 | SIGLEC1 | -3.48 | 6.559E-15 |
| GGT5 | 5.24 | 2.5976E-16 | MYO7A | -3.27 | 5.6658E-14 |
| ATP1B2 | 5.07 | 4.7851E-16 | MERTK | -2.96 | 4.6628E-10 |
| CASP5 | 5.03 | 5.019E-16 | RARRES1 | -2.67 | 1.2986E-05 |
| STAC | 4.41 | 8.996E-13 | IL10 | -2.60 | 1.4385E-06 |
| IRAG1 | 4.17 | 1.6605E-21 | SDS | -2.45 | 5.2349E-16 |
| CD177 | 4.01 | 8.9685E-11 | LGMN | -2.43 | 2.0277E-05 |
| PROK2 | 3.90 | 5.8345E-34 | SAMD4A | -2.31 | 6.1006E-10 |
| HSD11B1 | 3.65 | 1.6172E-09 | FUCA1 | -2.26 | 4.5911E-10 |
| LRG1 | 3.59 | 4.4558E-51 | ALDH1A1 | -2.26 | 7.2881E-13 |
| ALPL | 3.55 | 1.5464E-14 | L1CAM | -2.14 | 2.2154E-06 |
| FFAR3 | 3.49 | 5.1243E-27 | C2 | -2.12 | 1.4225E-06 |
| NAIPP1 | 3.45 | 8.7279E-08 | TLR7 | -2.05 | 6.6752E-09 |
| CCM2L | 3.44 | 4.9446E-09 | LINC00900 | -2.01 | 7.1447E-06 |
| CCL24 | 3.44 | 2.4222E-25 | CRISP3 | -1.89 | 0.00010407 |
| MOB3B | 3.34 | 2.3564E-32 | MS4A4A | -1.73 | 0.00086178 |
| CXCL5 | 3.24 | 6.7662E-11 | L3MBTL4 | -1.65 | 1.3195E-05 |
| FFAR1 | 3.23 | 6.5507E-14 | BAALC | -1.61 | 6.0227E-06 |
| SPATC1 | 3.20 | 1.6894E-22 | CMKLR1 | -1.58 | 1.062E-05 |
| SLPI | 3.19 | 1.3158E-09 | TREM2 | -1.56 | 9.0553E-07 |
| CSF1 | 3.11 | 2.902E-13 | KRT72 | -1.51 | 0.00047091 |
| ANXA3 | 3.03 | 1.1412E-05 | TSPAN4 | -1.50 | 3.1871E-06 |
| GALNT14 | 2.98 | 3.7614E-07 | GJB2 | -1.46 | 1.1631E-05 |
| PLIN5 | 2.93 | 4.6215E-12 | ARMC9 | -1.46 | 1.3581E-06 |
| MMP3 | 2.90 | 2.0059E-06 | STARD13 | -1.45 | 0.00010416 |
| IL1RAP | 2.89 | 3.2708E-43 | ADAP2 | -1.45 | 1.3826E-05 |
| CD22 | 2.81 | 9.8078E-29 | GPNMB | -1.32 | 0.00027575 |
| CLEC4A | 2.77 | 1.8591E-33 | P2RY11 | -1.30 | 9.4504E-07 |
| SLC1A2 | 2.73 | 4.0097E-06 | STAB1 | -1.29 | 0.00067141 |
| OLIG2 | 2.72 | 5.2177E-06 | EPB41L3 | -1.28 | 2.6028E-06 |
| CSF3 | 2.71 | 0.00681077 | MARCHF1 | -1.22 | 3.5403E-08 |
| GSEC | 2.68 | 6.8308E-14 | RASGRP3 | -1.21 | 1.3844E-05 |
| FAM157A | 2.65 | 6.6461E-24 | C2CD2 | -1.18 | 5.4366E-05 |
| FCER2 | 2.64 | 3.7443E-12 | ADAMDEC1 | -1.18 | 0.00039694 |
| DCN | 2.64 | 0.00477356 | LIPA | -1.18 | 2.0689E-05 |
| MRC1 | 2.61 | 5.6989E-13 | ADA | -1.17 | 3.8608E-08 |
| TREML3P | 2.57 | 3.2618E-27 | ADIRF-AS1 | -1.14 | 8.5748E-05 |
| LINC01094 | 2.56 | 8.0024E-25 | SCARB2 | -1.13 | 1.0144E-09 |
| PI3 | 2.55 | 7.6446E-05 | TCN2 | -1.11 | 0.00017738 |
| FAM157B | 2.53 | 2.4191E-09 | SLC2A8 | -1.09 | 3.255E-06 |
| TEAD3 | 2.52 | 7.8273E-07 | LACC1 | -1.09 | 2.1692E-06 |
| MCEMP1 | 2.45 | 9.1922E-06 | SLC12A7 | -1.08 | 1.0713E-05 |
| SLC25A37 | 2.41 | 1.3285E-22 | NR1H3 | -1.07 | 8.5861E-05 |
| MACC1 | 2.39 | 0.00340973 | PTGIR | -1.07 | 3.2699E-05 |
| SLC25A27 | 2.39 | 1.31E-11 | ARHGAP22 | -1.06 | 3.7329E-06 |
| CISH | 2.38 | 1.0789E-28 | CYFIP1 | -1.04 | 0.00070611 |
| CR1 | 2.36 | 3.7316E-07 | IL4I1 | -1.03 | 2.9538E-05 |
| SOCS3 | 2.36 | 6.6074E-24 | HSH2D | -1.01 | 9.3973E-07 |
| CXCL1 | 2.36 | 6.6074E-24 | ALDH2 | -1.01 | 5.5905E-05 |

Computed using DESeq2. Significance was calculated using the Wald test (two-tailed) followed by the Benjamini-Hochberg correction

[a] Differential gene expression considered significant when adj. p-value cutoff < $10^{-4}$

Log2FC; Log2 fold change, Adj. p-value; adjusted p-value

### Supplemental Table 10

**Supplemental Table 10** Top 50 genes up- and downregulated in ADSC cultured with iMSC-conditioned neutrophils compared to ADSC cultured with MSC-conditioned neutrophils

| Gene name | Average Log2FC | Adj. p-value[a] | Gene name | Average Log2FC | Adj. p-value[a] |
|---|---|---|---|---|---|
| **Up when cultured with iMSC-cond. neutrophils** | | | **Down when cultured with iMSC-cond. neutrophils** | | |
| FCER2 | 5.68 | 7.76E-19 | RGS4 | -1.27 | 1.12E-19 |
| CSF3 | 4.08 | 3.55E-09 | TNFRSF11B | -1.14 | 1.44E-05 |
| SERPINB4 | 3.88 | 7.68E-05 | ASPN | -0.86 | 8.56E-08 |
| CCL24 | 3.31 | 1.91E-05 | CD36 | -0.84 | 8.00E-09 |
| MRC1 | 2.70 | 8.50E-17 | DKK1 | -0.74 | 1.67E-05 |
| IL24 | 2.68 | 1.26E-17 | CRYAB | -0.67 | 7.15E-07 |
| CXCL8 | 2.61 | 1.50E-10 | LRRC2 | -0.64 | 0.000974 |
| MCEMP1 | 2.60 | 8.27E-09 | | | |
| CXCL1 | 2.59 | 3.42E-26 | | | |
| CXCL5 | 2.29 | 8.69E-10 | | | |
| FBP1 | 2.28 | 8.81E-10 | | | |
| IL36RN | 2.27 | 7.69E-07 | | | |
| ITGAM | 2.21 | 1.56E-09 | | | |
| CXCL6 | 2.21 | 1.84E-18 | | | |
| CXCL2 | 2.09 | 7.21E-09 | | | |
| MMP8 | 2.04 | 2.63E-09 | | | |
| HSD11B1 | 1.97 | 8.12E-05 | | | |
| IL1B | 1.97 | 0.00011 | | | |
| SPN | 1.83 | 2.41E-05 | | | |
| STEAP4 | 1.83 | 4.61E-05 | | | |
| PTGS2 | 1.81 | 0.000335 | | | |
| CYBB | 1.75 | 3.71E-17 | | | |
| TDO2 | 1.66 | 2.87E-10 | | | |
| IL3RA | 1.64 | 0.000229 | | | |
| CCRL2 | 1.61 | 4.00E-06 | | | |
| IL6 | 1.61 | 1.30E-06 | | | |
| CD300LF | 1.60 | 0.000261 | | | |
| NFKBIZ | 1.60 | 5.84E-08 | | | |
| JAML | 1.52 | 4.57E-06 | | | |
| CLEC12A | 1.51 | 2.22E-05 | | | |
| C3 | 1.50 | 1.67E-05 | | | |
| IL1RN | 1.46 | 6.27E-05 | | | |
| LILRB3 | 1.43 | 1.94E-07 | | | |
| ITGAX | 1.43 | 2.21E-07 | | | |
| LIF | 1.42 | 7.93E-05 | | | |
| TXNIP | 1.39 | 9.48E-09 | | | |
| PTAFR | 1.38 | 4.92E-06 | | | |
| ZC3H12A | 1.38 | 1.50E-10 | | | |
| IGSF6 | 1.35 | 0.000113 | | | |
| CFB | 1.35 | 5.51E-11 | | | |
| HK3 | 1.32 | 0.000405 | | | |
| CD22 | 1.29 | 1.27E-05 | | | |
| AQP9 | 1.29 | 3.83E-11 | | | |
| APLN | 1.26 | 1.56E-09 | | | |
| SERPINA1 | 1.25 | 1.67E-05 | | | |
| PDPN | 1.24 | 2.08E-09 | | | |
| CCR1 | 1.14 | 8.11E-07 | | | |
| CXCR4 | 1.12 | 0.000491 | | | |
| VLDLR | 1.09 | 6.82E-05 | | | |
| CD53 | 1.09 | 0.000465 | | | |

Computed using DESeq2. Significance was calculated using the Wald test (two-tailed) followed by the Benjamini-Hochberg correction

[a] Differential gene expression considered significant when adj. p-value cutoff < $10^{-3}$

Log2FC; Log2 fold change, Adj. p-value; adjusted p-value

### Supplemental Table 11

**Supplemental Table 11** Genes up- and downregulated in ADSC cultured with iMSC-conditioned neutrophils in the presence of an isotype control compared to in the presence of anti-IL-1β.

| Gene name | Average Log2FC | Adj. p-value[a] | Gene name | Average Log2FC | Adj. p-value[a] |
|---|---|---|---|---|---|
| **Up when cultured in the presence of isotype control** | | | **Down when cultured in the presence of isotype control** | | |
| IGHG4 | -2.55 | 2.24E-12 | FAT3 | 1.68 | 0.001657 |
| CXCL6 | -1.46 | 2.03E-08 | ZNF329 | 1.48 | 0.000216 |
| CXCL3 | -1.27 | 1.96E-09 | HMGCS1 | 0.77 | 2.51E-10 |
| CXCL1 | -1.26 | 1.26E-11 | CEMIP | 0.45 | 0.000255 |
| CXCL2 | -1.15 | 1.26E-11 | APP | 0.42 | 0.00156 |
| CXCL8 | -1.08 | 1.07E-18 | SCD | 0.41 | 0.000451 |
| CXCL5 | -1.07 | 1.49E-09 | HSPB7 | 0.40 | 2.73E-05 |
| IL1B | -0.88 | 1.49E-09 | FADS2 | 0.25 | 0.002337 |
| IL1RN | -0.74 | 6.58E-09 | FAT3 | 1.68 | 0.001657 |
| KYNU | -0.65 | 0.00084 | ZNF329 | 1.48 | 0.000216 |
| MMP9 | -0.60 | 0.006477 | HMGCS1 | 0.77 | 2.51E-10 |
| MMP3 | -0.55 | 4.65E-05 | CEMIP | 0.45 | 0.000255 |
| ITGB2 | -0.50 | 0.000255 | APP | 0.42 | 0.00156 |
| SERPINB2 | -0.50 | 0.000793 | SCD | 0.41 | 0.000451 |
| PTGS2 | -0.48 | 0.004759 | HSPB7 | 0.40 | 2.73E-05 |
| MMP1 | -0.41 | 0.001597 | FADS2 | 0.25 | 0.002337 |
| TNFRSF1B | -0.39 | 0.007299 | | | |
| PLAU | -0.26 | 0.008279 | | | |

Computed using DESeq2. Significance was calculated using the Wald test (two-tailed) followed by the Benjamini-Hochberg correction

[a] Differential gene expression considered significant when adj. p-value cutoff < $10^{-3}$

Log2FC; Log2 fold change, Adj. p-value; adjusted p-value

### Supplemental Table 12

**Supplemental Table 12** Top 50 differentially expressed genes (DEGs) comparing clusters MSC1 – MSC5 of treated MM to that of controls

| Gene name | Average Log2FC | Adj. p-value[a] | Gene name | Average Log2FC | Adj. p-value[a] |
|---|---|---|---|---|---|
| **Up in MM versus controls** | | | **Down in MM versus controls** | | |
| FABP4 | 0.631 | 9.44E-269 | FRMD6-AS1 | -0.340 | 0 |
| FABP5 | 0.620 | 0 | RP11-431K24.1 | -0.207 | 0 |
| CXCL2 | 0.534 | 5.72E-202 | RP11-182L21.6 | -0.222 | 2.10E-304 |
| BCYRN1 | 0.496 | 2.85E-142 | LRRC4C | -0.230 | 3.54E-280 |
| APOD | 0.482 | 2.10E-254 | RP11-758H9.2 | -0.322 | 1.70E-180 |
| CXCL8 | 0.391 | 1.71E-74 | RP11-47A8.5 | -0.324 | 5.34E-180 |
| EGR1 | 0.384 | 4.26E-41 | WDR74 | -0.292 | 1.13E-175 |
| JUNB | 0.379 | 2.01E-81 | RP11-796E2.4 | -0.219 | 2.63E-173 |
| TDO2 | 0.376 | 4.42E-296 | LINC00936 | -0.201 | 1.96E-158 |
| ADAMTS1 | 0.369 | 2.57E-12 | HP | -0.413 | 8.25E-144 |
| C10orf10 | 0.366 | 8.89E-09 | HIST1H4C | -0.295 | 1.47E-121 |
| IFI27 | 0.363 | 5.96E-247 | RP11-386I14.4 | -0.319 | 2.61E-105 |
| ADM | 0.360 | 6.02E-43 | HIST1H2AC | -0.203 | 7.60E-99 |
| SFRP1 | 0.360 | 1.26E-131 | PTCH2 | -0.240 | 1.10E-87 |
| ZFP36 | 0.342 | 2.50E-48 | TLR4 | -0.327 | 1.04E-86 |
| SOCS3 | 0.336 | 1.20E-61 | KCNQ1OT1 | -0.262 | 5.35E-74 |
| TGFBI | 0.328 | 3.23E-189 | KCNE4 | -0.376 | 2.12E-68 |
| CCL2 | 0.328 | 2.49E-16 | MROH8 | -0.409 | 2.06E-60 |
| JUN | 0.325 | 1.17E-77 | EFNA1 | -0.206 | 6.93E-57 |
| RGS16 | 0.323 | 5.41E-246 | CRH | -0.327 | 1.66E-55 |
| FOS | 0.317 | 3.56E-63 | HBB | -0.228 | 3.56E-53 |
| RRAD | 0.314 | 1.37E-240 | PPP1R10 | -0.214 | 2.14E-34 |
| PRRX1 | 0.312 | 0 | RP11-473M20.16 | -0.315 | 2.21E-32 |
| TRIB1 | 0.306 | 1.54E-50 | RNU12 | -0.250 | 4.82E-31 |
| DUSP1 | 0.302 | 1.97E-67 | CYB5D2 | -0.633 | 8.51E-20 |
| HLA-DRB5 | 0.298 | 0 | HNRNPUL1 | -0.266 | 4.86E-19 |
| ACTA2 | 0.297 | 0 | TNFAIP2 | -0.208 | 4.97E-17 |
| CHI3L2 | 0.287 | 2.05E-144 | SPEG | -0.218 | 5.01E-08 |
| NFKBIA | 0.279 | 2.63E-12 | WNT4 | -0.325 | 3.39E-05 |
| HLA-DRB1 | 0.279 | 6.15E-89 | | | |
| TIMP3 | 0.275 | 3.44E-16 | | | |
| FOSB | 0.275 | 2.28E-60 | | | |
| IGFBP5 | 0.272 | 9.95E-19 | | | |
| NCALD | 0.272 | 0 | | | |
| IGFBP3 | 0.263 | 0.000109 | | | |
| AHR | 0.262 | 1.77E-151 | | | |
| NNMT | 0.261 | 4.06E-15 | | | |
| VASP | 0.260 | 0 | | | |
| KLF2 | 0.244 | 9.82E-15 | | | |
| KLF4 | 0.243 | 4.86E-87 | | | |
| PPM1N | 0.239 | 0 | | | |
| PTX3 | 0.234 | 6.06E-143 | | | |
| HLA-DPA1 | 0.229 | 3.26E-31 | | | |
| MYC | 0.221 | 1.79E-102 | | | |
| CITED2 | 0.221 | 3.77E-25 | | | |
| CD14 | 0.219 | 5.09E-62 | | | |
| CYR61 | 0.217 | 6.43E-43 | | | |
| CH25H | 0.214 | 0 | | | |
| MAP1B | 0.213 | 0 | | | |
| NEAT1 | 0.213 | 5.96E-17 | | | |

Computed using Seurat's FindMarkers() function

[a] Differential gene expression considered significant when adj. p-value cutoff < $10^{-4}$

Log2FC; Log2 fold change, Adj. p-value; adjusted p-value

### Supplemental Table 13

**Supplemental Table 13** Top 50 differentially expressed genes (DEGs) comparing cluster MatNeu2a-b of treated MM to that of controls

| Gene name | Average Log2FC | Adj. p-value[a] | Gene name | Average Log2FC | Adj. p-value[a] |
|---|---|---|---|---|---|
| **Up in MM versus controls** | | | **Down in MM versus controls** | | |
| HIST1H1C | 0.125 | 4.70E-93 | MMP9 | -0.295 | 1.52E-31 |
| CXCL8 | 0.124 | 4.61E-25 | RPL26 | -0.215 | 3.97E-18 |
| SOD2 | 0.118 | 2.92E-14 | RPL6 | -0.189 | 2.21E-55 |
| PTGS2 | 0.112 | 7.25E-22 | IL1RN | -0.171 | 9.87E-61 |
| HMGN2 | 0.105 | 3.09E-55 | HIST1H2AC | -0.149 | 1.19E-29 |
| ANKRD49 | 0.104 | 1.27E-129 | PGLYRP1 | -0.142 | 1.92E-33 |
| IGSF6 | 0.099 | 2.20E-57 | RPL23A | -0.139 | 0.000779 |
| B3GNT5 | 0.099 | 1.33E-11 | CAMP | -0.115 | 2.54E-83 |
| NFKBIA | 0.082 | 3.30E-13 | PLBD1 | -0.111 | 9.53E-49 |
| SULT1B1 | 0.080 | 7.17E-56 | PPIF | -0.108 | 1.32E-47 |
| SET | 0.077 | 6.68E-92 | GADD45B | -0.107 | 1.34E-08 |
| H1FX | 0.076 | 1.45E-85 | RPL13A | -0.102 | 4.74E-12 |
| CREB5 | 0.074 | 6.36E-23 | ANXA3 | -0.102 | 1.51E-198 |
| G0S2 | 0.074 | 8.59E-06 | RPS8 | -0.098 | 3.13E-14 |
| ANP32E | 0.070 | 1.26E-23 | RETN | -0.090 | 4.22E-164 |
| SVIP | 0.069 | 1.08E-80 | TBCA | -0.088 | 1.37E-24 |
| SAT1 | 0.068 | 5.84E-08 | CD83 | -0.080 | 1.93E-18 |
| NEAT1 | 0.067 | 2.17E-26 | LUCAT1 | -0.079 | 7.42E-76 |
| MX2 | 0.064 | 5.56E-37 | ADM | -0.078 | 3.70E-195 |
| IL1RAP | 0.063 | 1.80E-17 | LTF | -0.077 | 1.26E-08 |
| TMEM123 | 0.059 | 2.79E-39 | PHACTR1 | -0.076 | 0.000379 |
| ANKRD28 | 0.058 | 0.000786 | DIS3 | -0.074 | 9.74E-07 |
| RNF213 | 0.058 | 4.84E-41 | LCN2 | -0.074 | 3.31E-60 |
| RIPK2 | 0.056 | 3.87E-78 | IFIT3 | -0.071 | 3.02E-135 |
| MT-ND4L | 0.055 | 5.33E-47 | RPS21 | -0.070 | 7.35E-09 |
| RP11-1143G9.4 | 0.049 | 7.83E-60 | PRPF4B | -0.067 | 7.28E-13 |
| CTD-3252C9.4 | 0.046 | 2.13E-52 | JUN | -0.066 | 3.12E-06 |
| GBP2 | 0.046 | 8.14E-51 | EIF3E | -0.064 | 1.35E-35 |
| STOM | 0.045 | 8.07E-05 | CD52 | -0.063 | 6.02E-20 |
| NCL | 0.041 | 2.58E-30 | SNHG8 | -0.063 | 3.06E-08 |
| SRGN | 0.041 | 2.59E-14 | CD24 | -0.062 | 8.59E-08 |
| SLC8A1 | 0.041 | 2.78E-22 | EEF1A1 | -0.060 | 0.000166 |
| IFITM1 | 0.040 | 8.39E-20 | IFIT2 | -0.058 | 1.34E-56 |
| IER5 | 0.040 | 3.68E-34 | MAFB | -0.057 | 2.42E-12 |
| FAM76A | 0.038 | 6.81E-98 | SMC4 | -0.057 | 9.22E-74 |
| AHNAK | 0.038 | 1.19E-58 | MTDH | -0.057 | 6.37E-27 |
| SMC1A | 0.035 | 2.60E-12 | HP | -0.056 | 3.02E-200 |
| SREK1IP1 | 0.035 | 3.69E-26 | IFI6 | -0.056 | 7.93E-05 |
| OTUD1 | 0.035 | 4.83E-98 | MMP8 | -0.055 | 4.88E-84 |
| RPSA | 0.035 | 2.43E-43 | UGP2 | -0.055 | 1.31E-88 |
| EEA1 | 0.033 | 2.22E-44 | SYNE1 | -0.054 | 3.57E-36 |
| RP13-942N8.1 | 0.033 | 2.72E-10 | TNFAIP2 | -0.053 | 1.80E-44 |
| CHI3L1 | 0.033 | 1.66E-51 | NDUFB2 | -0.053 | 1.16E-72 |
| FFAR2 | 0.033 | 2.74E-108 | MCM7 | -0.052 | 1.73E-44 |
| RPL18A | 0.032 | 8.15E-06 | ICAM1 | -0.052 | 1.04E-37 |
| KIF22 | 0.032 | 6.49E-44 | GJB6 | -0.052 | 5.29E-58 |
| MZT2B | 0.031 | 1.49E-30 | DEFA3 | -0.052 | 1.69E-157 |
| SERBP1 | 0.030 | 5.62E-21 | RPS20 | -0.051 | 4.04E-33 |
| FCGR1A | 0.030 | 9.60E-87 | TIMP1 | -0.050 | 2.55E-61 |
| TNFAIP6 | 0.030 | 1.22E-62 | GPR155 | -0.049 | 1.77E-123 |

Computed using Seurat’s FindMarkers() function

[a] Differential gene expression considered significant when adj. p-value cutoff < $10^{-4}$

Log2FC; Log2 fold change, Adj. p-value; adjusted p-value

### Supplemental Table 14

**Supplemental Table 14** Baseline characteristics of patients and non-myeloma controls included in proteomic analyses

| Variable | Myeloma | Non-myeloma controls |
|---|---|---|
| **N** | 52 | 17 |
| **Age** [a] | | |
| *Mean (y)* | 56 | 62 |
| *Range (y)* | 36- 65 | 26-80 |
| **Sex** | | |
| *Female (%)* | 42 | 29 |
| *Male (%)* | 58 | 71 |
| **R-ISS** | | N/A |
| *1 (%)* | 46 | |
| *2 (%)* | 42 | |
| *3 (%)* | 12 | |
| **Risk stratification** | | N/A |
| *Standard risk (%)* | 92 | |
| *High risk (%)* | 8 | |
| **Bone marrow plasma cells** [b] | | N/A |
| *Mean (%)* | 50 | |
| *Range (%)* | 10-100 | |
| **rFLC** | | N/A |
| *Mean* | 242 | |
| *Range* | 0-3080 | |
| **Plasma M-protein** | | N/A |
| *Mean (g/dL)* | 3.2 | |
| *Range (g/dL)* | 0-9.6 | |
| **MRD after treatment** [c] | | N/A |
| *Negative (%)* | 73 | |
| *Positive (%)* | 27 | |
| **Best confirmed therapy response**[d] | | N/A |
| *sCR (%)* | 24 | |
| *CR (%)* | 11 | |
| *VGPR (%)* | 57 | |
| *PR (%)* | 8 | |

[a] at time of tissue collection
[b] as determined by CD38+/CD138+ flow cytometry in bone marrow aspirates
[c] determined by flow cytometry, cut-off at $10^{-5}$
[d] as graded and reported by investigators
(y) years, sCR; stringent complete response, CR; complete response, VGPR; very good partial response, PR; partial response
